# Pervasive historical introgression across oak phylogeny shapes gene expression

**DOI:** 10.64898/2026.07.16.739029

**Authors:** Yuxiang Zhu, Ruirui Fu, Mingming Zhang, Yao Li, Xinyue Teng, Jing Wu, Ying Liu, Yuqing Zhang, Yan Lin, Zeyuan Zhang, Chunmei Pang, Hanxing Wu, Antoine Kremer, Martin Lascoux, Jun Chen

## Abstract

Oaks were once described as the “worst-case scenario” for the biological species concept because of the presumed pervasive hybridization and introgression among species. However, we still lack a full picture of genome-wide introgression across the eight sections of the *Quercus* phylogeny. Furthermore, we still do not have a complete characterization of what is introgressed and how it functionally influences the recipient species. Firstly, we used 168 whole genomes from 67 *Quercus* species belonging to the eight sections of the *Quercus* phylogeny and a super pan-genome to generate a phylogeny and test for ILS and introgression. Introgression was widespread: ancient introgression occurred between ancestors of the eight sections and more recent introgression took place within sections. Secondly, using genome-wide transcriptome, methylome, and chromatin accessibility data from a subset of 19 species sampled from four sections we demonstrated that introgressed genes are more highly and uniformly expressed than non-introgressed ones. This difference in expression level between introgressed and non-introgressed genes mostly results from *cis*-regulation, and is most likely due to DNA methylation decay in promoter ACRs. In summary, introgression has been a major evolutionary force during the whole evolutionary history of *Quercus* and had a significant impact on fundamental processes such as gene regulation and expression.

## Introduction

Since Darwin’s time, most biologists have accepted that a species is hard to define but, generally speaking, consists of a group of reproducing individuals that are, somehow, reproductively isolated from other individuals and can be defined by a set of specific traits (Coyne & Orr, 2004; Mallet, 2004). As Darwin himself wrote “No one definition has as yet satisfied all naturalists; yet every naturalist knows vaguely what he means when he speaks of species” (Darwin, 1859). For instance, most botanists and foresters can easily agree on what differentiates a pedunculate from a sessile oak, even if most of them are also keenly aware that the two species are not strictly reproductively isolated and that hybrids are indeed quite common (Leroy *et al*., 2020). However, what most biologists of the 20^th^ century could probably not have foreseen is the sheer amount of gene exchange that did and, in many cases, still does take place, among what appears as well-defined species. The last decades have indeed shown that hybridization and introgression in natural populations were much more widespread than anticipated (Edelman & Mallet, 2021) even though, in most genera, the true extent of hybridization and introgression still remains to be estimated. Introgression, on which we are going to focus here, describes the transfer of genetic material between related species through hybridization and backcrossing to the parental species (Aguillon *et al*., 2022).

Considering that introgression appears to be much more widespread than anticipated, a first question is to what extent introgression contributes to the evolution of classical phylogenetic subdivisions, such as lineages and sections. Does introgression occur across the entire phylogeny or is it restricted to sections? Although evolutionary categories such as sections are still hard to define, they nonetheless indicate some degree of evolutionary integrity in either morphology or genotype that would be expected to limit or prevent gene flow. In general, hybridization and introgression do not appear to be uniformly distributed across species. In plants, for instance, Mitchell *et al*. (2019) showed that woody perennial species tend to hybridize more than annuals or herbaceous species. Estimates of the frequency of hybridization and introgression were often obtained from pairs of closely related species but introgression could be equally pervasive in ancestral and modern lineages. Introgression may even be more extensive across ancestral lineages since crossing barriers are likely to be weaker between recently diverged species. Hence, studies on a higher phylogenetic subdivision level, e.g., a genus or a family, other than closely related species, should be carried out to fully understand the extent of the evolutionary influence of introgression (Suvorov *et al*., 2022; Cicconardi *et al*., 2023). In trees, previous studies suggest that intersectional gene flow did occur in both oaks (Crowl *et al*., 2020; Hipp *et al*., 2026) and Eucalypts (Orel *et al*., 2026).

A second, and related, question is whether introgressed genes differ in their mode and level of expression between the donor and recipient species. Surprisingly, while there have been numerous studies testing whether introgressed fragments are adaptive (reviewed in (Hedrick, 2013; Suarez-Gonzalez *et al*., 2018)), little attention has been paid to the mechanisms by which introgressed alleles can alter the phenotypic variation of the recipient species that sometimes diverged millions of years ago. For instance, is this achieved through protein coding sequence changes or through modifications in gene expression regulation (McCoy *et al*., 2017; Silvert *et al*., 2019)? Are introgressed genes expressed differently from non-introgressed ones? Are introgressed genes expressed differently across species? Or, on the contrary, do we observe convergent evolution of gene expression at introgressed genes? If they are differentially expressed, is the change in level of expression due to changes in *cis* or in *trans* regulation?

The genus *Quercus*, commonly known as oaks, is an ideal system for studying introgression since it includes more than 400 species in the Northern Hemisphere, most of which are sympatric or parapatric with at least one other oak species (**Fig. 1A**). Evidence of hybridization and introgression has been reported at both morphological and genetic level in both closely and distantly related species (Leroy *et al*., 2020; O’Donnell *et al*., 2021; Zhou *et al*., 2022). In our previous study on two sympatric Asian oaks, we detected enriched signals of adaptive introgression in transposable elements that can serve as *cis*-regulatory elements that affect the gene expression in the two species (Fu *et al*., 2022). However, in oaks and numerous plant species, hybridization and introgression are seldom limited to pairs of species and can be pervasive through the whole evolutionary history of the genus (e.g., (Hipp *et al*., 2020)). In the present study, we aimed to address the following questions.

**Figure 1.**
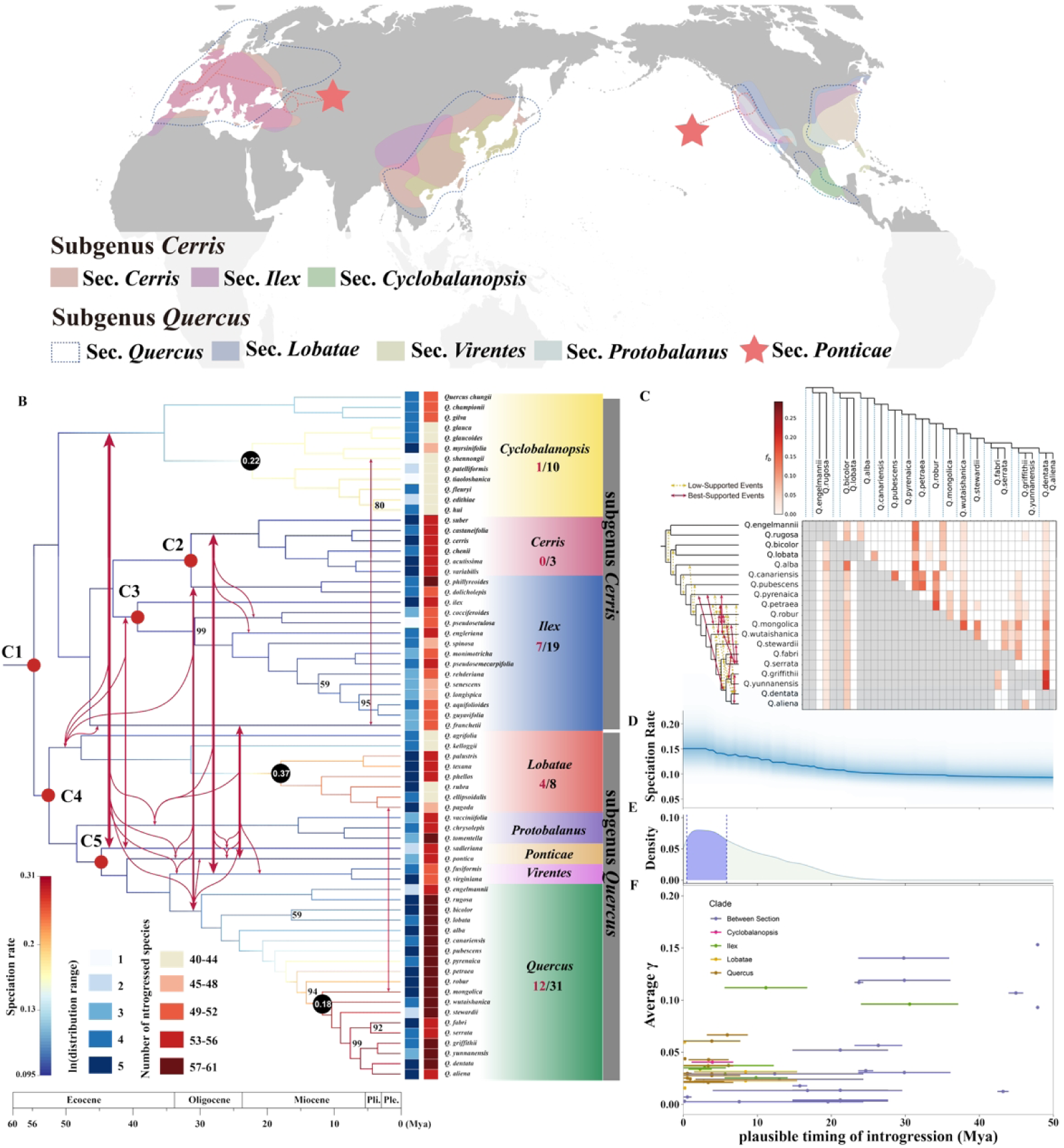
The distribution ranges, phylogenetic tree, and introgression events inferred among 67 oak species. **(A)** The distribution ranges for the oak species included in this study; **(B)** The phylogenetic tree was inferred based on 168 genomes of 67 oak species aligned to a super-pan genome of the genus *Quercus*. The branches were colored by the lineage speciation rate. The number in each black circle at the three nodes gives the posterior probability that a shift in diversification rate occurred at that node. All supporting bootstrapping values are equal to 100 except those specified at the nodes. The red arrows illustrated the best-supported inter-sectional introgression events. The black number below each section name shows the total number of intra-sectional introgression events while the red shows the number of best-support events; The distribution range is given on a log scale. Log(distribution range) presents the number of 1°×1° grids in log-scale. **(C)** An example of *f*-branch test in section *Quercus*. Arrows indicate the intra-sectional introgression events of best-supported (red) or low-supported (yellow); **(D)** The distribution of speciation rates; **(E)** The distribution of TE insertion times; **(F)** The time and fraction of introgressed genome for the 58 best-supported introgression events.

Firstly, how widespread is introgression across the oak phylogeny? Is introgression restricted within sections or does it also occur among sections? We explore occurrences of introgression throughout the genus *Quercus* by sequencing the whole genome of 168 individuals coming from 67 species and belonging to the 8 botanical sections of the genus (Hipp *et al*., 2020). In order to facilitate the assembly and comparisons of those genomes we created a super-pan genome using nine oak species. We then used an array of methods to estimate the number of introgression events within and between sections and the fraction of the genome affected by introgression. Secondly, we compared introgressed and non-introgressed genes and asked whether they are differently expressed and whether some biological functions are overrepresented among introgressed genes. Finally, using a subset of 77 individuals from 19 oak species representative of the different sections we compared gene expression and its regulation (*trans*-*vs. cis*-regulatory elements) between introgressed and non-introgressed genes using transcriptome (RNA-seq), methylome (WGBS), and chromatin accessibility data (ATAC-seq).

## Results

### The phylogeny inferred from the super pan-genome of Quercus

A super pan-genome of the *Quercus* genus was generated using nine high-quality genomes at the chromosomal level (**Fig. S1**) and a ML phylogenetic tree with high statistical support was inferred based on whole genome sequences of 168 individuals from 67 species (**Fig. 1 & Fig. S2**). As in previous oak phylogenetic studies (e.g. Denk *et al*., 2017; Hipp *et al*., 2020; Yang *et al*., 2021; Zhou *et al*., 2022), all species could be assigned to two subgenus and eight sections. While other species in *Ilex* are monophyletic, *Q. phillyreoides*, *Q. dolicholepis*, and *Q. franchetii*, that also belonged to section *Ilex* in a previous study (Hipp *et al*., 2020), were grouped in section *Cerris* or in the basal position of section *Ilex* and *Cerris*. As *Ilex* and *Cerris* are two closely related sections, these differences with previous clustering are likely due to incomplete lineage sorting (ILS) and/or introgression. To be conservative and consistent with previous studies, we filtered out the conflicting topologies involving these three oak species when identifying introgression events. The divergence times between sections are in general agreement with those obtained by (Hipp *et al*., 2020; Zhou *et al*., 2022).

### Introgression is widespread across the Quercus phylogeny and not restricted to sections

An identity-by-descent (IBD) analysis revealed long shared haplotype blocks in a highly structured pattern within each phylogenetic section (**Fig. S3**), suggesting extensive gene flow within section. We then calculated the *f*_d_ statistic to test for the existence of introgression events in all possible species triplets using *Castanopsis delavayi* as an outgroup. In total, 66.5% of the triplets (33,314/50,117), covering 78.8% of all possible species pairs (1,743/2,211), had a *f*_d_ statistic value that significantly deviates from zero, of which 1,486 (85.3%) are between species pairs from different sections and 257 (14.7%) are between species pairs from the same section (**Fig. S5C & D**). Rescaled by the total number of species pairs within each section, on average 84.9% of species pairs within section *Quercus* may have experienced introgression, followed by 75.2%, 69.2%, and 61.5% of species pairs experiencing introgression within section *Ilex*, *Lobatae*, and *Cerris*, respectively. For introgression between species from different sections, on average 83.3% of all species pairs tested had significant *f*_d_ statistic (**Fig. S5D**). Species from either *Quercus* or *Cerris* had the highest probability of intersectional gene flow (>90%) while species from *Lobatae* had the lowest (62.9%, **Fig. S5D**).

To estimate the most parsimonious number of introgression events across the phylogeny and the time at which introgression took place, we first calculated a heuristic matrix of *f*-branch values. We then assigned introgression events to the most recent common ancestor of a group of species if introgression signals were detected in more than 80% of its descendants. In addition, we also tested for introgression by using two complementary tests: a Discordant Count Test (DCT) and a Branch Length Test (BLT). These two tests allow to differentiate patterns of discordance due to introgression from those due to ILS alone (Suvorov *et al*., 2022). The best-supported introgression events were kept only if the two branches that experienced introgression overlapped in the dated phylogenetic tree. In the end, we identified 45 best-supported introgression events and 21 of them occurred between species of different sections (**Fig. 1B & C** and **Fig. S4**). The proportion of introgressed genome () was plotted against the time when introgression events took place (γ varied from 0.25% to 15.3% with a median equal to 3.1%, **Fig. 1F**). For introgression events between sections, 14 out of 21 took place above 20 Mya. Introgression events within sections were much more recent, on average, with 16 out of 24 having their midpoint time between 0.16 and 3.9 Mya. Only two introgression events in section *Ilex* occurred before 10 Mya (11.16 and 30.58 Mya). For both inter-and intra-sectional introgression events, the proportion of introgressed genome, γ, was significantly correlated with the estimated time at which introgression occurred (*p*-value = 1.87e-3 and 1.10e-3 for within and between section introgression events, respectively; *p*-value = 1.32e-4 for combined). Interestingly, the timing of increased speciation rate coincided with that of TE expansion and intra-sectional introgression events (**Fig. 1D-F**).

Introgression was widespread both between and within sections, albeit with some variation. On average, we found that each oak species introgressed with 40 to 61 other oak species. However, species from the section *Quercus* introgressed with the largest number of species of all sections (mean = 58.1), while species from sections *Lobatae* and *Cyclobalanopsis* introgressed with the lowest number of species (mean = 46.8 and 44.3, respectively). After correction for phylogeny, species with larger distribution ranges introgressed with more species (*p*-value = 0.037) than those with more restricted ranges. Species that introgressed with a larger number of species also tended to have slower lineage speciation rates (*p*-value = 0.00229) with the exception of the section *Quercus*, where Asian and non-Asian species had very similar introgression levels but significantly different speciation rates (0.3 for Asian and 0.11 for non-Asian oaks, respectively). Introgression and the extent of overlap distribution areas were not significantly correlated.

### Introgressed genes are common, highly pleiotropic and associated with reproduction

Genomes of all oak species were divided into windows of 10 kb long and the *f*_d_ statistic was calculated in each genomic window for all triplets. The *f*_d_ statistic was significant in 19,988 windows. Of all 30,249 genes, 11,769 were in windows with a significant *f*_d_ statistic, of which 10,676 introgressed between species across all eight sections. Only five genes introgressed between species of a single section (**Fig. 2A**). The *f*_d_ statistic increased with the number of sections that a gene was introgressed into and, on average, one gene was found to be introgressed among 63 oak species (**Fig. 2B**). To better capture the introgression dynamics of genes, we calculated the gene introgression frequency, which measures the probability that one gene is introgressed between two randomly picked species in all triplets. The introgression frequency increased with GC3 content and the recombination rate (Spearman’s *p*-value = 7.3 × 10^-4^ and 3.1 × 10^-13^, **Fig. 2C and D**). Furthermore, 11,589 genes were introgressed between more than two species (mean introgressed species number = 64.1, 95% CI: 28 – 68) and only 21 genes were introgressed between only two species, both with strong supports (>90% of bootstraps). These two categories are, hereafter, referred to as “widely-introgressed genes” and “specifically-introgressed genes”, respectively. 159 genes were introgressed between more than two species but with weak support (<90% of bootstraps, mean introgressed species number = 16.9, 95% CI: 3 – 68). Those genes are referred to as “ambiguously-introgressed genes” hereafter. See definitions in **M&M**). The widely-introgressed genes had significantly higher introgression frequency than ambiguously-introgressed and specifically-introgressed ones (mean = 7.11% versus 0.83% and 0.04%, **Fig. 2E**).

**Figure 2.**
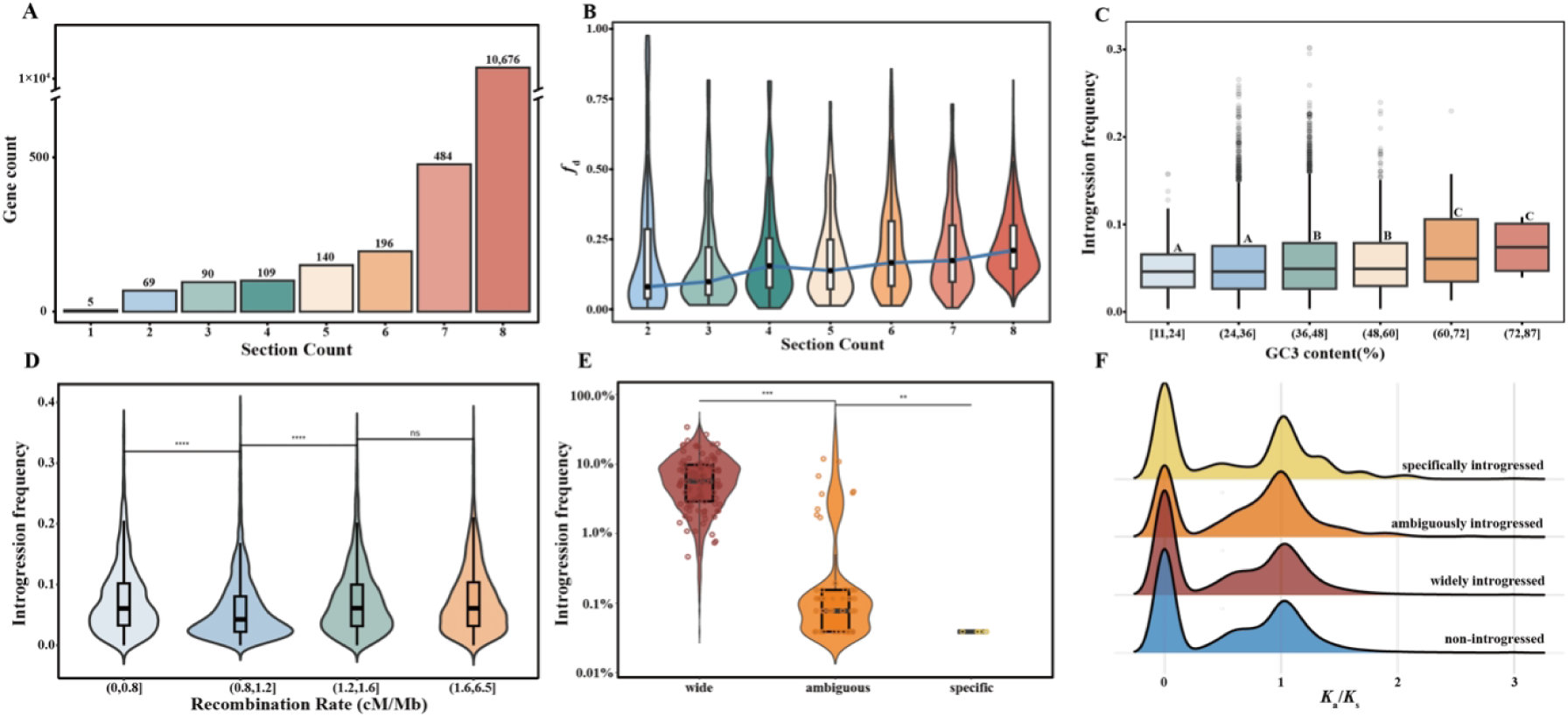
The gene count, introgression frequency and *Ka*/*Ks* distributions for introgressed genes across eight phylogenetic sections. The number of introgressed genes **(A)** and *f*_d_ value **(B)** across the eight different sections; The introgression frequency increases with GC3 content (C) and recombination rate (D); **(E)** The introgression frequency for genes that introgress among multiple species with high-confidence (“widely introgressed”), among multiple species but with low-confidence (“ambiguously introgressed”), and strictly between two species (“specifically introgressed”); **(F)** the distribution of *Ka*/*Ks* for four groups of genes.

No significant difference was found between the *K_a_*/*K_s_* distributions of introgressed and non-introgressed genes (**Fig. 2F**). A protein-protein interaction analysis indicated that introgressed genes had on average slightly higher pleiotropic degree than non-introgressed ones (mean = 54.9 versus 51.0, *p*-value = 0.001) (**Fig. 3A**). In addition, a plant ontology (PO) enrichment analysis showed that introgressed genes were expressed in multiple tissues, and, more importantly, were enriched in genes coding for floral organs (**Fig. 3B**). The gene ontology (GO) enrichment analysis also confirmed that introgressed genes were functionally enriched for reproductive processes, like ovule and pollen development, as well as other processes related to stress response and cell growth (**Fig. 3C**). No significant signals in PO or GO enrichment analyses were found for non-introgressed genes.

**Figure 3.**
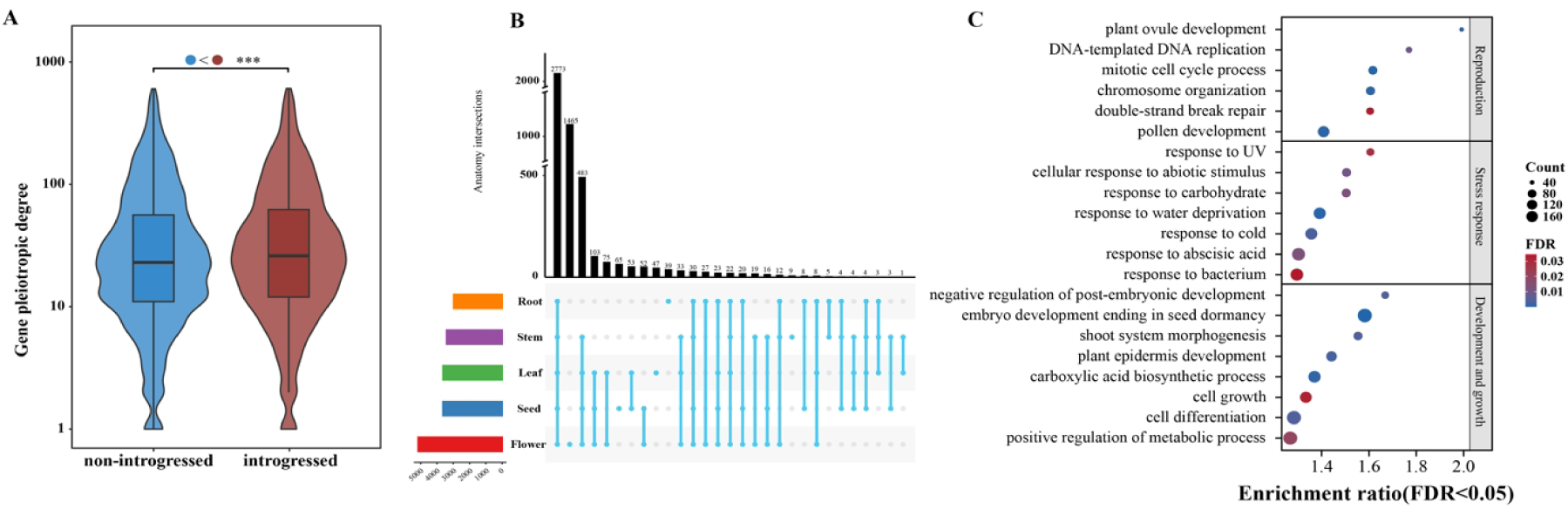
The pleiotropy of introgressed genes. **(A**) Gene pleiotropic degrees for introgressed and non-introgressed genes; **(B)** Number of tissues where the expression of introgressed genes was found based on Plant Ontology analysis; **(C)** Putatively related biological processes of introgressed genes based on Gene Ontology enrichment analysis.

### The methylation and chromatin accessibility explain the high and conserved expression of introgressed genes across species

The gene expression, methylation, and accessibility of chromatin regions (ACRs) were studied on a subset of 77 individuals from 19 oak species representative of four different sections (see **Table S1**). Introgressed genes were more highly expressed and their expression level was more conserved across species than non-introgressed ones (median log (TPM+1) = 2.34 and 0.6, median CV = 1.02 and 1.51, respectively; *p*-values < 2.2e-16, **Fig. 4A - C**). Such patterns remained when controlling for the difference in gene number between the two groups (**Fig. S6**). To explore the regulatory mechanisms of gene expression, we compared the expression regulatory mechanisms between introgressed and non-introgressed genes. This was done by measuring *cis*-regulation by the number of ACRs in the promoter and the methylation level along the gene body and promoter regions, and *trans*-regulation by the number of transcription factors binding to the target gene.

**Figure 4.**
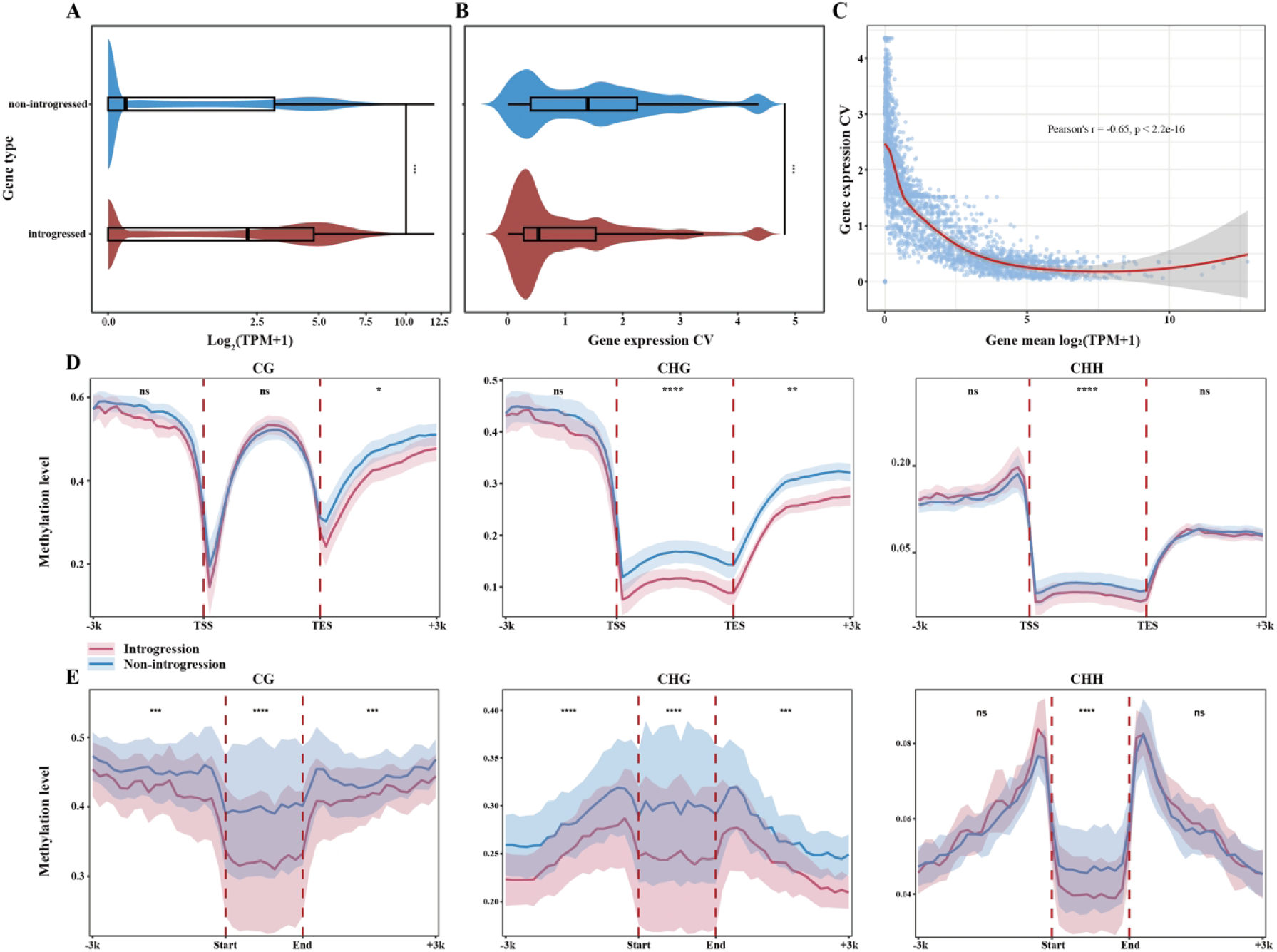
The different expression and methylation profiles for introgressed and non-introgressed genes. **(A)** and **(B)** The distributions of gene expression level and coefficient of variance of expression across species for introgressed and non-introgressed genes; **(C)** gene expression coefficient of variation (CV) decreases when gene expression level increases; **(D)** The distributions of methylation level for introgressed and non-introgressed protein coding genes and **(E)** for ACR regions. CG, CHG, and CHH refer to the three main contexts of DNA methylation in plants, where the C is a cytosine, and the G (guanine) and H (A, C, or T) are adjacent nucleotides. TSS and TES indicate a transcription start and end site for protein coding genes **(D)** and the start and end indicate the start and end of ACR **(E)**. The flanking area is 3 kb each. For *p*-value of significance: ** <0.05, *** <0.001, **** < 0.0001, n.s.: > 0.05.

Genes that have promoter ACRs and lower methylation level tend to be associated with higher expression levels (**Fig. S7**). Overall, 95.7% and 90.3% of introgressed and non-introgressed genes were methylated, respectively (**Table S11**). Introgressed genes had significantly lower methylation level in the genic region (without up-/downstream flanking regions) at the CHG and CHH sites (mean = 0.103 versus 0.153, *p*-value = 8e-14; mean = 0.023 versus 0.028, *p*-value = 3.1e-10, respectively), and in the 3kb downstream region at CG and CHG sites (0.405 versus 0.453, *p*-value = 0.03; mean = 0.234 versus 0.285, *p*-value = 0.002, respectively). Although CG dinucleotides had higher methylation levels than CHG and CHH for both groups of genes, a significant difference was only observed in the 3kb downstream region with lower methylation level for introgressed genes (**Fig. 4D and Table S12**).

The presence or absence of promoter ACRs for each gene was compared between introgressed and non-introgressed genes using logistically transformed values and a generalized linear model with binomial distribution. Introgressed genes are more likely to harbor one or more promoter ACRs than non-introgressed genes (65.8% versus 47.2%, *p*-value < 2.2e-16) and the positions of ACRs in introgressed genes are also closer to the gene body (median = 940 bp versus 1043 bp). For the genes containing ACRs, the methylation levels of the genic region and the promoter ACRs were also compared for both introgressed and non-introgressed genes. Almost all genes containing ACRs (99.8% and 99.76% for introgressed and non-introgressed genes, respectively) had their genic regions methylated and the methylation level was significantly lower in introgressed genes (mean mCG = 0.417 versus 0.426, *p*-values = 9.536e-05, mean mCHG = 0.098 versus 0.146, *p*-values < 2.2e-16, mean mCHH = 0.022 versus 0.026, *p*-values < 2.2e-16). The methylation level of the promoter ACRs was also lower for introgressed genes (mean mCG = 0.32 versus 0.397, *p*-values < 2.2e-16, mean mCHG = 0.246 versus 0.3, *p*-values = 2.9e-16, mean mCHH = 0.04 versus 0.047, *p*-values = 5.4e-11; **Fig. 4E**). We also observed lower methylation levels in 3kb up-and downstream flanking regions for introgressed genes (see details with 95% CI in **Table S13**).

The percentages of transposable elements in promoter ACRs also differed between introgressed and non-introgressed genes. Seventeen oak species, out of 19, had significantly higher percentages of TE in promoter ACRs for non-introgressed genes (1.07 ∼ 1.69-fold, **Fig. 5, Table S14**). Moreover, promoter ACRs that contain TE had much higher methylation levels than those that do not contain TEs (1.57 ∼ 20.92-fold, *p*-value = 5.13e-12 for CG, 1.66 ∼ 18.89-fold, *p*-value = 2.87e-12 for CHG, and 1.79 ∼ 8.42-fold, *p*-value = 2.50e-10 for CHH. **Fig. 5**).

**Figure 5.**
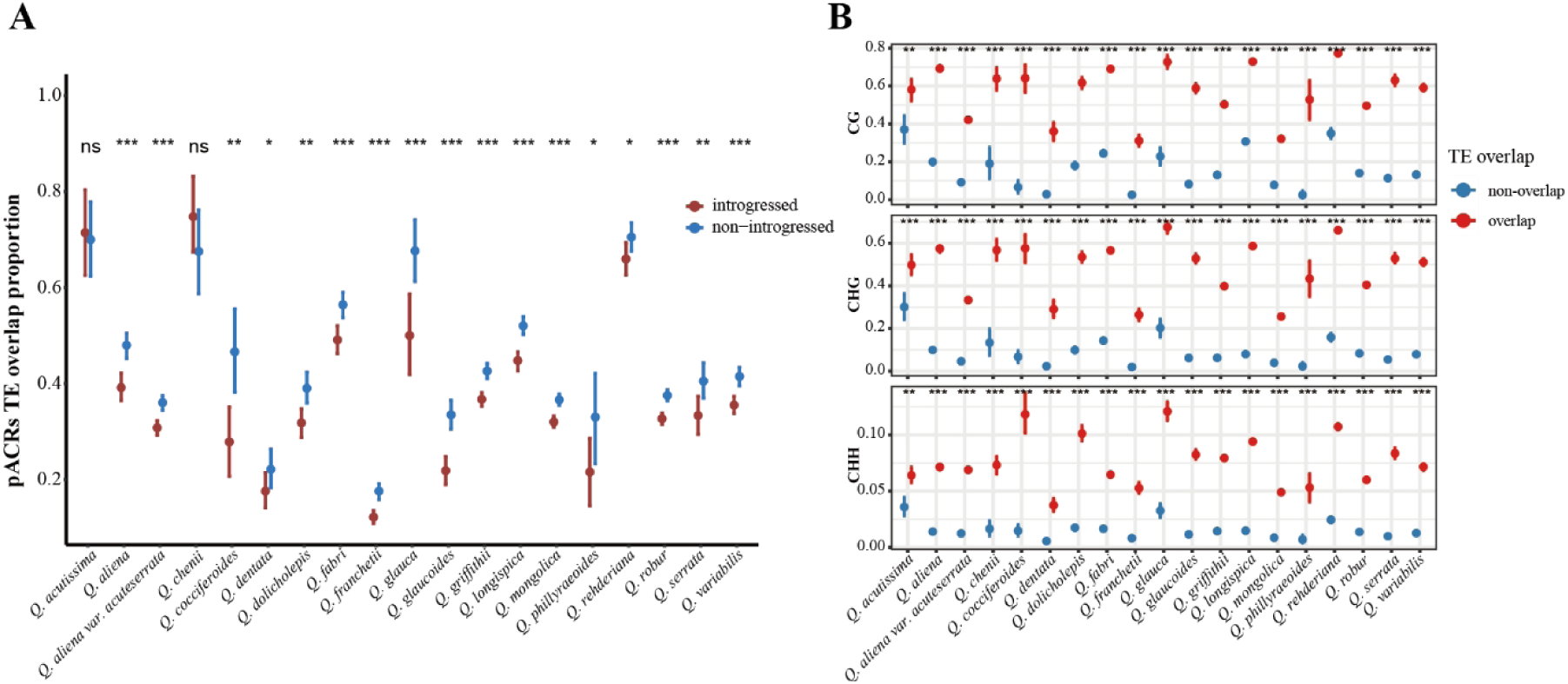
Transposable elements and methylation. (**A**) The percentages of promoter ACR containing TEs for introgressed and non-introgressed genes across the different species. (**B**) The methylation level for promoter ACRs with or without TE. Mean and 95% CI generated by bootstrapping were shown for each species. For p-value of significance: * <0.05, ** <0.01, *** <0.001, **** < 0.0001, n.s.: > 0.05.

### Introgressed transcription factors were not responsible for expression differences

Finally, regarding *trans*-regulatory elements, in total 1,200 transcription factor coding genes were identified and 621 of them harbored introgression signals, significantly more than what can be expected by chance (Hypergeometric test *p*-value = 1.49e-20). The proportion of introgressed genes varied across TF families. For instance, TF families NF-YA, HB-PHD, ZF-HD, LSD, and EIL include only introgressed genes, whereas families Whirly, VOZ, LFY, NZZ/SPL, S1Fa-like, and STAT include only non-introgressed genes. Family B3, on the other hand, had significantly fewer introgressed than non-introgressed genes (27 out of 75, *p*-value = 0.0034). The 57 families with MYB and bHLH contained significantly more introgressed genes (78 out of 119, *p*-value=4.87e-5; and 60 out of 96, *p*-value =0.0103, respectively) than other families. In general, introgressed TF genes also had higher gene expression levels and more conserved expression profiles across species. In particular, the seven introgressed TF enriched families also had more conserved expression profiles across species (**Fig. 6**).

**Figure 6.**
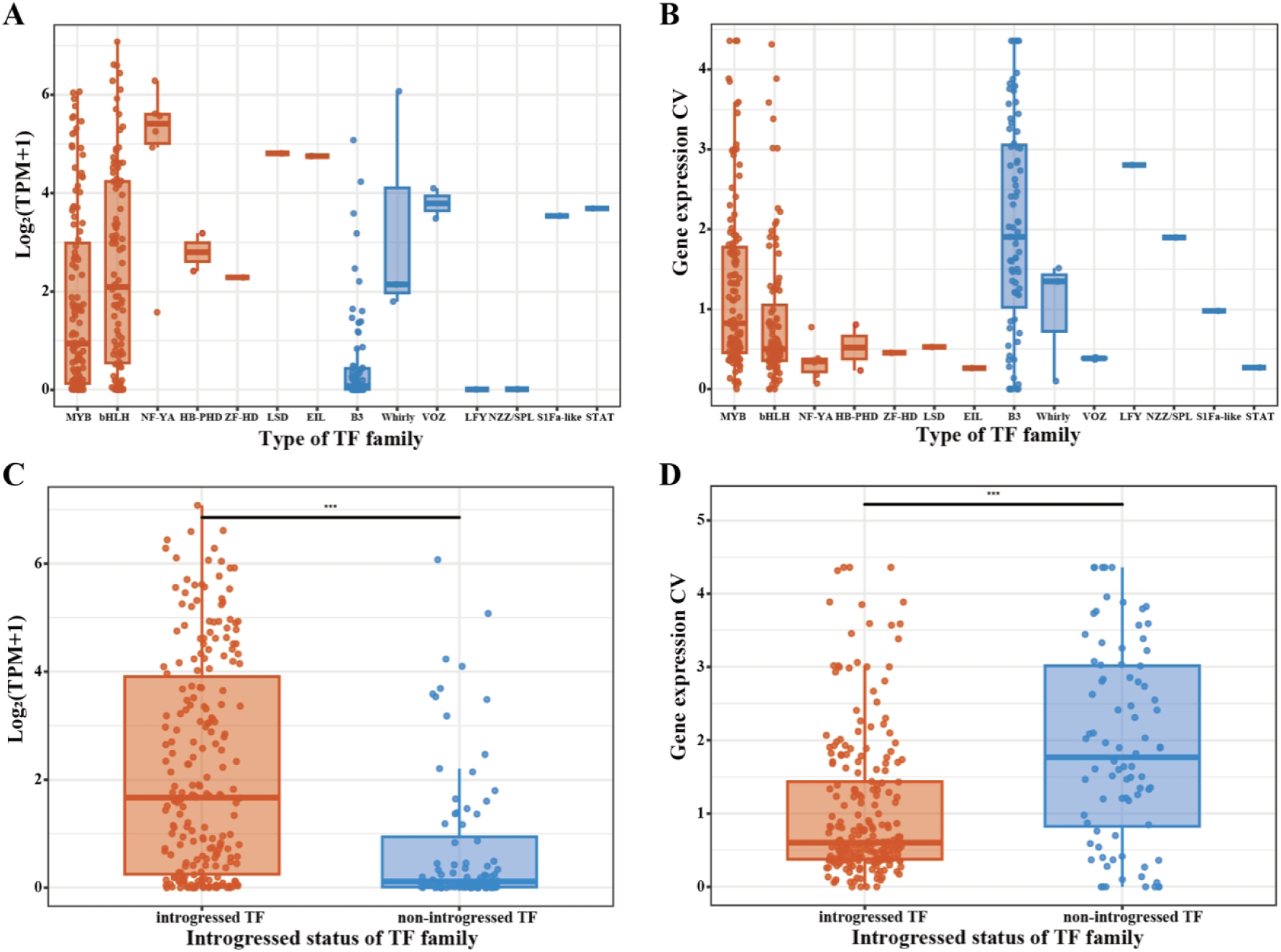
The distributions of gene expression and coefficient of variance across species for introgressed and non-introgressed genes that encode for transcription factors. **(A)** and **(B)** for the TF families mainly composed of introgressed genes (orange) or non-introgressed genes (blue). **(C)** and **(D)** for all introgressed (orange) and non-introgressed TF genes (blue).

We further explored whether introgressed genes are more frequent in downstream target genes. First, a total of 24,474 genes containing transcription factor binding sites were found. 12,044 genes only contained binding sites for introgressed TF genes while 142 genes only contained binding sites for non-introgressed TF genes. The remaining 12,288 genes contained binding sites for both types of TF genes. So, 621 introgressed TF genes, accounting for 51.8% of total TF genes, affect the expression of almost all downstream target genes (99.4%). 9,836 of the downstream target genes were introgressed genes and each on average harbored 29.4 TF binding sites with 26 of them binding to introgressed TF genes. 14,638 were non-introgressed genes that each contained on average 28 transcription factor binding sites with 24.6 binding to the introgressed TF genes. Hence, almost all downstream genes were affected by introgressed TF, and introgressed genes did not dominate among downstream genes.

In summary, patterns of both *cis*-and *trans*-regulatory elements differ strongly between introgressed and non-introgressed genes. However, marginal differences were detected in numbers of transcription factor binding sites between the two groups of genes. Hence the difference in expression level and pattern between introgressed and non-introgressed genes mostly resulted from *cis*-regulation. To quantify the relative contributions of phylogeny, introgression, and methylation level of *cis*-regulatory elements to gene expression, we used a Phylogenetic Generalized Linear Mixed model using either the gene expression level or its coefficient of variation as the response variable, and introgression frequency, methylation ratio at genic region and at the promoter ACRs as the fixed effects, and phylogeny as a random effect. Of all the variables with a significant effect (*p*-values < 5e-4), the introgression frequency had the largest positive effect on gene expression level (standardized effect size = 2.426, **Fig. 7, Table S15 & S16**). Methylation ratio at the genic region had the strongest negative effect on gene expression (standardized effect size = −10.227), followed by the methylation ratio at the promoter ACR (standardized effect size = −1.993). On the opposite, methylation ratio at the genic region had the strongest positive effect on the coefficient of variation of gene expression (standardized effect size = 2.54) and methylation at the promoter ACR had much smaller but significant effect (standardized effect size = 0.14), while introgression frequency had negative effects (standardized effect size = −0.32).

**Figure 7.**
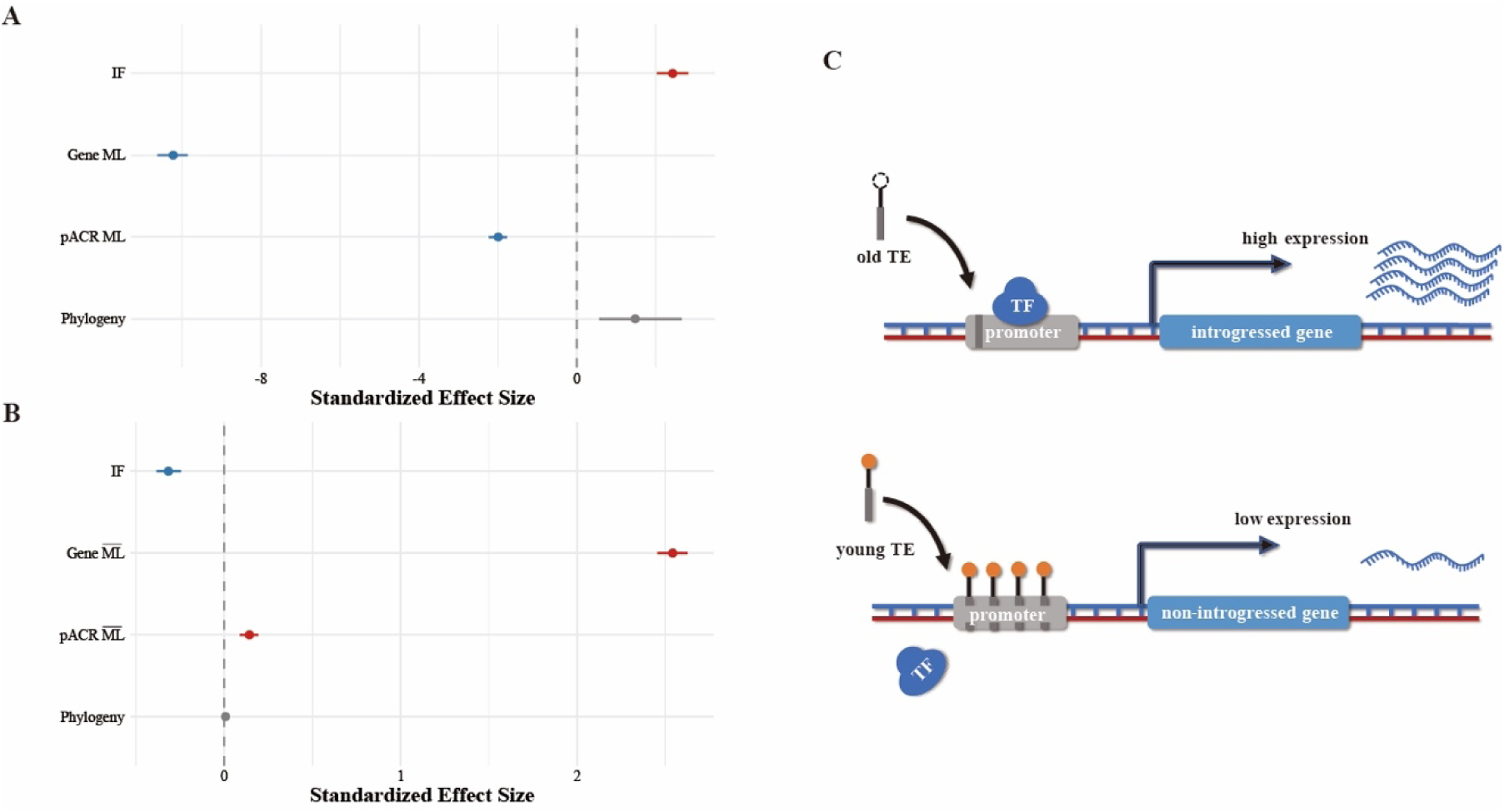
The standardized effect sizes for all variables affecting gene expression levels (A) and coefficient of variation (B). The red dots represent the positive effects while the blue dots represent the negative effect. The effect of phylogeny was considered as a random effect and illustrated in the gray triangle. IF: introgression frequency; ML: methylation level.

## Discussion

### The distribution of introgression in the genus Quercus

In the present study we leveraged extensive genomic, transcriptomic and epigenetic data and showed that introgression across the whole *Quercus* genus is extensive and occurred in a non-random pattern along the genome. Our study constitutes therefore a first step towards a deeper understanding of the mechanisms of introgression. First, we showed that introgression was not limited to within sections. Sections are not simply dividing a complex phylogenetic topology into monophyletic substructures, but they also represent groups of species with common morphological or life history traits characteristics. For instance, the ‘red oaks’ of sect. *Lobatae* have soft bristles on their leaves’ lobes, while ‘white oaks’ of sect. *Quercus* don’t. In oaks, however, section is never a barrier for gene flow (e.g., Crowl *et al*., 2020; Hipp *et al*., 2026). On the contrary, a large proportion of introgression events were identified between species belonging to different sections. In particular, introgression was found in almost all species pairs between a few sections, e.g., between sections *Ilex* and *Quercus*, *Cyclobalanopsis* and *Lobatae*, and is, paradoxically, relatively less common between any two species within section. Many of the species that experienced introgression do not exhibit overlapping distributions today, or are even found on different continents (e.g. introgression was identified between *Quercus ilex* in the Mediterranean and *Q. variabilis* in East Asia). Although long-distance dispersal can occur in species with wind-dispersed pollen, the critical dispersal distance of viable pollen grains for pollination is nonetheless generally limited to 25 ∼ 100 km (Schueler & Schlünzen, 2006). Hence, it is more likely that the signals of extensive intersectional introgression reflect historical gene flow between the ancestors of extant species and took place during the early stages of species divergence. Indeed, 21 ancestral introgression events were identified between sections using a parsimony method combining *f*-branch, DCT and BLT tests. Though not frequent, such inter-sectional gene flow was also reported between ancestors of the Eurasian *Roburoid* and *Q. pontica* and between North American *Dumosae* and *Prinoideae* lineages (Crowl *et al*., 2020), and was also found between two Chinese oaks belonging to different sections, *Q. fabri* (section *Quercus*) and *Q. acutissima* (section *Cerris*) (Li *et al*., 2021). The dearth of previous studies observing intersectional gene flow is likely due to the targeted focus on closely related and sympatric species when investigating hybridization and introgression, e.g., for section *Quercus* (Ribicoff *et al*., 2025), *Virentes* (Eaton *et al*., 2015) and *Lobatae* (Sullivan *et al*., 2016). Recently, based on genomic sequences of 54 oak species, Zhou *et al*. (2022) also identified intersectional introgression events, albeit much fewer than in the present study and mainly between sections *Quercus* and *Lobatae* and between sections *Quercus* and *Ponticae*. This discrepancy could be caused by differences in alignment methods used by the two studies. Zhou *et al*. (2022) aligned reads to a single reference genome of *Q. robur*. Alignment to a single reference genome significantly limits the mapping ratio of species from a different section. In contrast, we used a super pan-genome based on nine high quality genomes from the five largest sections of genus *Quercus* and the average mapping ratio was elevated to 89.14% (95% CI: 74.01 – 98.12%). A small part of the phylogenetic topology (i.e., section *Ilex*) based on SNPs identified in alignment to the super pan-genome differed from previous studies. Such conflicts, however, could not be caused by aligning to the super pan-genome as the same topology can also be fully recovered by a species tree based on 9,804 gene trees (**Fig. S8**). Nevertheless, those controversial uncertain branches were removed to ensure that the estimates of intersectional introgression events are reliable. It is also expected that with the increase of the number of species studied, more ancient introgression events at the early stage of species diversification will be discovered as was observed in many model organisms like *Heliconius* (Cicconardi *et al*., 2023) and *Drosophila* (Suvorov *et al*., 2022), as well as in the present study.

Secondly, it is not surprising to see that the estimated proportion of introgressed genome, γ, decays with the timing of introgression, or that *f*_d_ statistic decreases with species divergence, i.e., the branch length or *D*_xy_. Most intrasectional introgression events occurred about 5∼10 Mya when the species diversification rate also increased. While diversification accelerated, the number of species potentially undergoing introgression increased and the number of introgressed genomic regions tended to decrease with growing divergence. Our results further emphasize that historical gene flow had a profound effect on genome evolution, and shed light on introgression dynamics in extant species. Finally, our study showed that TE expansion was associated to introgression, as the majority of introgressed genomic windows contain TE. It would be interesting to explore the possibility that introgression facilitated the spread of TE across the *Quercus* genus, which further accelerated species diversification (Plomion *et al*., 2018) and, in turn, limited introgression.

Our results emphasize that introgression can act as a cohesive force within the genus, maintaining evolutionary potential. Introgressions can be seen here as the genetic component of the sympatric parallel adaptive radiation that prevailed throughout history and continues to shape community assemblages in oak ecosystems to this day (Cavender-Bares, 2019). In most places of the Northern hemisphere, historical biogeography indicates that the genus radiated by maintaining species belonging to different sections, for example *Quercus* and *Lobatae* in North America or *Cerris* and *Ilex* in Eurasia. Our results suggest that intersectional introgression was rather the driver than the consequence of the parallel radiation, by maintaining the potential for exploring a variety of ecological opportunities during colonization. Ultimately, this may not be the whole story as in a very few cases introgression failed to maintain the sectional assemblages, as for example when *Lobatae* went extinct in Europe and only *Quercus* expanded at the onset of the glacial-interglacial era. In summary, introgression in oaks is truly a genome-wide process, with up to one third of the protein coding genes introgressing between species of all sections, and no particular genomic region being enriched for introgression (**Fig. S9**).

### Potential adaptive role for introgression across the whole phylogeny

It may, at first glance, seem quite surprising that more than one third of the genes were introgressing between an average of 64 species and across all eight sections of the phylogeny. Based on the biology of oak species, one would rather expect introgressed genes to be under purifying selection and therefore rare. First, oak species usually have a relatively large effective population size, and deleterious mutations are expected to be purged very quickly and neutral mutations are not expected to be repeatedly observed across species. Second, introgressed genes also tend to be highly pleiotropic and involved in more protein-protein interactions than non-introgressed ones (Leroy *et al*., 2020; Fu *et al*., 2022). Frequent ancestral introgression between sections is enriched in functional processes like ovule, pollen, and embryo development, seed dormancy, and response to stress. These results suggest that intersectional introgressed could be adaptive and contribute to fast diversification between sections.

### The relationship between introgression and gene expression

In Fu *et al*. (2022) we showed that if individuals of *Q. acutissima* and *Q. variabilis* live in similar environments then introgression more frequently affect the same genes and introgressed genes exhibit similar expression profiles between the two species. The present study extends this result by revealing that introgressed genes have higher and more conserved expression than non-introgressed genes across almost all oak species between or within sections, again suggesting that introgressed genes may not evolve neutrally in oaks.

*Cis*-regulatory regulatory elements were the main drivers of gene expression differences between introgressed and non-introgressed genes in the case of Neanderthal-introgressed genes into modern humans (McCoy *et al*., 2017). The same appears to be true in oaks. Indeed, introgressed genes and their regulatory regions are more likely to be in ACRs and show lower methylation levels than non-introgressed ones. In plant promoter regions, all three methylation types, CG, CHG, and CHH are associated with reduced expression levels (Niederhuth *et al*., 2016). In the present study, the difference in methylation between introgressed and non-introgressed genes mainly occurred at the three types of methylation sites in the promoter ACRs, if present, and at CHH and CHG sites of the genic region, otherwise. Methylation in promoter regions prevents the binding of transcription activators and thereby favors the transcription repressors, thus leading to the inactivation or reduction of transcription. While methylation in genic regions inhibits the transcript elongation (Lucibelli *et al*., 2022), low methylation, on the contrary, helps to stabilize the expression of genes shared across different oak species. On the other hand, for *trans*-regulatory elements, even though higher proportions of TF genes were affected by introgression than the background genome (51.8% versus 38.9%), they had almost equal effects on all downstream target genes with little binding preference to introgressed or non-introgressed genes. Thus, the contribution of *trans* regulatory elements to expression differences between introgressed and non introgressed genes appears to be more limited than that of *cis* regulatory ones.

Transposable elements can also affect gene expression levels by acting as targets of DNA methylation in plants (Hirsch & Springer, 2017). In this study, we did not observe any significant difference in TE content in genic regions between introgressed and non-introgressed genes (67.9% and 67.8%) nor between the methylation levels of the TE (89.8% and 90.0%). In fact, it was the TE insertions in promoter ACRs that differed between the two groups of genes. In maize, ACR containing TEs can serve as gene promoter and facilitate gene amplification (Bubb *et al*., 2025). vonHoldt *et al*. (2012) showed that recent transposed LTR-RTs have a characteristic chromatin accessibility and the methylation pattern of LTR-RTs degenerates with evolutionary age. Thus, more recently inserted LTR-RTs had higher methylation levels. A possible explanation for the difference of TE content and DNA methylation in promoter ACRs could be that the TE insertions were rather recent in oak genomes and more likely affected non-introgressed genes and more recent and lineage specific introgression (**Fig. S10**).

The relationship between foreign DNA and gene expression regulation has been studied for many years though most studies were based on recombinant inbred lines developed by hybridizing natural and wild populations. For example, introgressed alleles from *Zinzania latifolia* Griseb into cultivated rice lines can also induced changes in DNA methylation and gene expression patterns (Liu *et al*., 2004). Rodriguez Barreto *et al*. (2019) also reported transgenerational effects of epigenetic introgression in Atlantic salmon. In modern humans, McCoy *et al*. (2017) showed that one quarter of Neanderthal-introgressed alleles exhibited *cis*-regulatory effects, that contributed to human phenotypic diversity and varied across tissues. A later study (Silvert *et al*., 2019) further showed that the regulatory effects were the consequence of introgressed variants in the promoter and enhancer regions, thereby suggesting underlying epigenetic changes. Furthermore, Hibbins and Hahn (2021) demonstrated that introgression can affect thousands of quantitative traits, including gene expression levels across the genome, and thus generate convergent patterns of evolution in the genus *Solanum*. Our study, for the first time, associated ancestral introgressed alleles to gene expression patterns and epigenetic regulatory elements across species of an entire genus. At this stage, this conclusion should be taken with a grain of salt since transcriptomic and epigenetic data were only available for 19 species, out of a very large genus of more than 400 to 5000 species. Nonetheless, our study provides a solid hypothesis about the genetic mechanisms by which introgression can affect the evolution of a large group of phylogenetically related species.

## General conclusion

In summary, our study demonstrated the presence of widespread introgression events across 67 oak species representatives of the genus *Quercus*. Ancient introgression occurred between sections, while more recent introgression took place within sections. Our study further revealed that introgressed genes are characterized by low methylation at promoter ACR and genic region leading to a significantly higher level of expression of introgressed genes than non-introgressed ones (**Fig. 7C**). Furthermore, the variance in expression level of introgressed genes across species is remarkably low. The high percentage of introgressed gene and their conserved expression provide the genetic basis for the pleiotropic functions of introgressed genes and contribute to the shared features of oaks. This high level of introgression across oak species also appears consistent with the general concept on an evolution on two fronts proposed by (Burger, 1975) cited in Hipp (2025) to solve the apparent paradox of the presence of “classical species” in the face of extensive gene flow and introgression. On a first front, at a more global level, all oak species are embedded within a single “syngameon” within which single species exchange genes. On a second front, the highly pleiotropic introgressed genes contribute to the adaptation of each species to its ecological niche, either directly or indirectly. Hence, a better characterization of introgressed genes could be an important step towards solving the apparent paradox of the maintenance of species in presence of extensive gene flow.

## Materials and Methods

### Plant materials & data collection

In this study, we constructed a comprehensive *Quercus* dataset consisting of 168 individuals from 67 species representing all eight recognized sections of the genus *Quercus*. Two individuals from *Lithocarpus dealbatus* and two individuals from *Castanopsis delavayi* were included as outgroup species. 83 re-sequenced individuals from 55 species were retrieved from the public domain while 89 re-sequenced individuals were produced in the current study (**Table S1**). Additionally, transcriptomic sequences (RNA-seq) and whole-genome bisulfite sequences (WBGS) were also produced for 77 individuals of 19 species while chromatin accessibility sequences (ATAC-seq) were produced for one individual in each of 19 species using fresh leaf tissues of one year old seedlings (**Table S1**). Acorns were germinated and grown in a growth chamber under a 12 h photoperiod, with temperature of 26°C, and a photosynthetic photon flux density (PPFD) of over 300 μmol m^−2^s^−1^ in Zhejiang University for one year. Fresh leaves were sent to Novogene company for sequencing.

### Genetic variants calling using Quercus super pan-genome

Genomic DNA was extracted from fresh leaves frozen in −80 ℃ using DNAsecure Plant Kit (Tiangen Biotech) according to the manufacturer’s protocol and paired-end short reads with a length of 150 bp were generated for each sample on Illumina platforms with PE150 strategy with a mean depth of 20×. A super pan-genome was generated using high-quality chromosome-level genomes of nine *Quercus* species downloaded from public databases, which represent five of eight sections in *Quercus* phylogeny (**Table S2.** Please refer to the **supplementary texts 1.2** for the constructure of super pan-genome). Sequencing adaptors were trimmed and cleaned short reads were aligned to the super pan-genome using software ‘vg giraffe’ that is implemented in vg v5.1.2 (Garrison *et al*., 2018; Siren *et al*., 2021) (**Table S3**). Alignment GAM files were converted to PACK files by ‘vg pack’ with parameters ‘-Q 10 -s 5’. The VCF files containing individual genetic variants were generated by ‘vg call’ based on PACK files and pan-genome GFA file. Then bcftools v1.13 (Danecek *et al*., 2021) was used to merge all VCF files, and low-quality SNPs were filtered out by vcftools with parameters ‘--max-missing 0.2 --minQ 30’. The final VCF file contained 9,795,402 SNPs.

### Phylogenetic inference

We constructed two maximum likelihood phylogenetic trees using IQ-TREE2 v.2.0.7 (Minh *et al*., 2020), one based on all 168 individual genomes and the other based on 67 species for which we randomly kept one individual genome for each species. *Castanopsis delavayi* was selected as an outgroup to root the phylogenetic tree. To minimize the effect of linkage disequilibrium, we performed SNP thinning with a 500 bp interval using VCFtools (--thin 500 parameter), which retained 537,748 SNPs. The best nucleotide substitution model and corresponding base frequency parameters were determined by Bayesian Information Criterion (BIC) model selection method implemented in IQ-TREE2. For both individual based and species based phylogenetic trees, the ‘TVM+F+R2’ model was chosen for the nucleotide substitution model, base frequency, and rate heterogeneity, respectively. Ultrafast bootstrap was run with 5,000 iterations to obtain the branch support for each node with parameter ‘-bb 5000 -bnni’.

### Identification of ancestral introgression events

We evaluated the percentage of individual species genome affected by introgression using *D*-statistic test and a modified *f*-statistic (*f*_d_) (Martin *et al*., 2015; Malinsky *et al*., 2021). To identify introgression events among all *Quercus* species, genome-wide ABBA-BABA test was conducted on 50,117 species triplets ((P1, P2), P3) using Dsuite Dtrios (MALINSKY et al., 2021) based on the species tree topology with *Castanopsis delavayi* as the outgroup (P4) (**Table S6**). Results were filtered by thresholds Z-score > 3 and false discovery rate (FDR) < 0.01. We switched order of species in ((P1, P2), P3) triplet and only kept significant combinations with *D*-statistic > 0 to test for introgression between P2 and P3. This resulted in a total of 38,993 triplets with significant introgression signals. Due to the bias of the *D*-statistic in small genomic regions, we also applied the *f*_d_ statistic to avoid inflated values in regions of reduced diversity (Martin *et al*., 2015). The *f*_d_ statistic was calculated using ‘calculate_abba_baba.r’ script (Martin *et al*., 2015) and was filtered with criteria 0 < *f*_d_ ≤ 1, Z-score > 3 and FDR < 0.01, which retained 33,314 high-confidence triplets with significant introgression signals. Finally, based on the *f*_d_ statistic we summarized introgression events between P2 and P3 in one phylogenetic section or between two sections by calculating the ‘introgression prevalence’, which is defined as the number of unique P2 and P3 combinations in significant triplets divided by the number of all possible P2 and P3 combinations across all species within tested sections.

To obtain an estimate of the most parsimonious number of introgression events across all *Quercus* species included in this study, we performed a two-step protocol on the results of the ABBA-BABA tests. Firstly, we performed a branch-specific introgression identification using the *f*-branch method implemented in Dsuite to distinguish introgression events between species tips from those occurring between ancestral branches. This approach calculates evolutionary lineage-resolved *f*-branch metrics that minimize spurious correlations while localizing introgression to specific phylogenetic branches. The analysis was conducted with a significance threshold of *p*-value < 0.05 followed by visualization of branch-level introgression signals using the ‘dtools.py’ script. Introgression events were assigned to the most recent common ancestor if *f*-branch values were greater than zero in more than 80% of its descendant species.

Secondly, two tests were carried out to identify introgression signals between branches identified in the first step. The first test is the ‘discordant-count’ test (DCT) and it is based on the number of gene trees that are discordant with the species tree. The second test is the branch-length test (BLT) and it relies on distorted coalescent time caused by introgression and/or ILS (Suvorov *et al*., 2022). A total of 9,804 gene trees were used in both analyses and each gene tree was constructed on windows of 1,000 SNPs using IQ-TREE2 with ‘GTR+I+G’ model (Dong *et al*., 2022). ASTRAL v1.16.3.4(Zhang *et al*., 2018) was used to reconstruct species tree based on 9,804 gene trees (**Fig. S8**). As for the *f*-branch test, at least 80% of descendant species in tested two ancestral branches were required for an introgression event to be identified by either DCT or BLT tests. A supporting score was calculated as the number of significant triplets in both DCT and BLT tests divided by the total number of triplets. Finally, we only kept introgression events with a supporting score larger than 10% and for which the time intervals of the two branches experiencing introgression overlap (**Table S7**). Additionally, we used the “blt_dct_test.r” script to estimate the introgression proportion γ across the genome for each introgression event. γ is defined by the following equation:

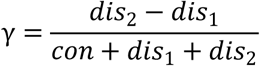

where *con* represents the number of concordant gene trees, and *dis_1_* and *dis_2_* represents the numbers of discordant gene trees of two possible topologies, and *dis_2_* > *dis_1_*.

### Sliding window introgression detection

To localize the introgression along the genomes for each pair of species where significant introgression had been identified, we divided genomes into non-overlapping sliding windows of 500bp and 10kb, respectively and calculated *D* and *f*_d_ in each window using the Python script ‘ABBABABAwindows. py’ (Martin *et al*., 2015). Each 10kb window was further divided into 20 blocks and the standard error of the *D*-statistic was estimated using the ‘leave-one-out’ jackknife approach, which was then used to obtain a normalized Z-score (Green *et al*., 2010; Zhang *et al*., 2016). *p*-value was calculated based on Z-score and multiple testing effect was controlled using the R package ‘qvalue’. The same approach was applied to calculate *f*_d_ and the significance *p-*values. Genomic windows experiencing significant introgression were selected as D > 0, 0 < *f*_d_ ≤ 1 and q-values of both *D* and *f*_d_ < 0.01.

To quantify how many oak species exchanged genetic variants for a given introgressed genomic windows, we calculated an ‘introgression frequency’ index with the following bootstrapping protocols: 1) in a given pair of species P2 and P3, where introgression had been identified with the genome-wide ABBA-BABA test, we chose a triplet ((P1, P2), P3) and examined if a significant introgression signal can be detected in a 10 kb window. 2) Since there could be more than one species available as P1 and the significance of introgression can be affected by the choice of P1, we randomly chose one P1 at a time and repeated with replacement for 1,000 iterations. 3) we repeated the above steps over all possible P2 and P3 pairs in the given genomic window. 4) We then summarized the total number of P2 and P3 pairs between which introgression had been detected in this window as the ‘introgression frequency’. Finally, we classified all introgressed windows into three categories: 1) ‘widely-introgressed’ windows in which more than one pairs of species exchange genetic variants with a bootstrap confidence of 95% or higher. 2) ‘specifically-introgressed’ windows in which only one pair of species had introgression with a bootstrap confidence of 95% or higher. And 3) the remaining introgressed windows, named ‘ambiguously introgressed’ windows, in which introgression was detected in one or more pairs but with a bootstrap confidence lower than 95%.

We took the mean *f*_d_ value across all unique P2 and P3 species as the introgression strength for the introgressed window. The introgression strength of genes within the window was calculated by weighting the mean *f*_d_ of the window by the gene length. For genes across more than one windows, gene *f*_d_ value was weighted by the fraction of gene overlapped with that window and summed across all overlapping windows. We also classified genes into three categories using the above protocols with an additional step that a gene was classified as ‘ambiguously introgressed’ if the gene category conflicted with the window category.

### RNA-seq expression differentiation analyses

We generated transcriptome (RNA-seq), whole genome methylation (WGBS), and chromatin accessibility (ATAC-seq) data on a subset of 77 individuals from 19 oak species belong to four sections (sect. *Cerris*, *Ilex*, *Cyclobalanopsis*, and *Quercus*) using frozen fresh leaves tissues. Two fully elongated leaves of the first flush were collected from three to seven one-year-old seedlings (except only two seedlings were collected for *Q. glaucoides*) and total RNA and genomic DNA were extracted for RNA-seq and WGBS. ATAC-seq sequencing was carried out only for one sample per species (**Table S1**). Reads were aligned to the corresponding closest reference genome of the six (*Q. acutissima*, *Q. aquifolioides*, *Q. dentata*, *Q. gilva*, *Q. mongolica*, *Q. variabilis*).

RNA-seq libraries were prepared using the TruSeq RNA Library Kit (Illumina) and sequenced on a NovaSeq 6000 platform with paired-end reads of 150 bp long. Sequencing adaptors were trimmed and reads with ambiguous nucleotide bases were removed before being aligned to reference genomes using GMAP and GSNAP (Wu *et al*., 2016) (**Table S8**). Only concordantly aligned reads with proper paired and unique mapping positions were kept for expression differentiation analyses. The number of reads overlapping with any position from the beginning to the end of each gene were counted by featureCounts (Liao *et al*., 2013). Read counts per gene were then normalized to Transcripts Per Kilobase Million (TPM). Pearson’s correlation coefficients were computed between biological replicates of RNA data (**Fig. S11**). All pairwise correlation coefficients between biological replicates exceeded 0.8 except two individuals from *Q. chenii* and *Q. glauca*. An average value across biological replicates was used to represent the gene expression level of a given species. Gene expression levels were compared between orthologous genes across all 19 oak species.

### Whole-genome bisulfite sequencing and methylation calling

Genomic DNA degradation and contamination was assessed by agarose gels. DNA purity was checked using NanoPhotometer® spectrophotometer (IMPLEN, CA, USA). After DNA quality control testing, a positive control DNA sample was added and then all samples were fragmented into 200-400bp using Covaris S220. Single-stranded DNA (ssDNA) fragments were bisulfite-treated using Accel-NGS Methyl-Seq DNA Library Kit (Illumina, Swift). For library preparation, fragmented DNA samples were subjected to methylation sequencing adapters ligation, size selection and PCR amplification. Library quality was assessed on the Agilent Bioanalyzer 2100 system and pair-end sequencing of sample was performed on an Illumina platform (Illumina, CA, USA). Sequencing adapters were trimmed using Trimmomatic (Bolger *et al*., 2014) and raw WGBS reads containing adapters, low-quality nucleotides and unrecognizable nucleotide (N) were filtered out using fastp v0.19.7 (Chen *et al*., 2018). The clean reads were aligned to reference genomes using BISCUIT v1.3.0 (Zhou *et al*., 2024) (**Table S9**). Methylated cytosines (mCs) were identified from the uniquely mapped reads using ‘pileup’ and ‘vcf2bed’ functions in BISCUIT with the following parameters ‘biscuit vcf2bed -e -t c’. Methylation ratios of cytosines covered by at least three reads were calculated as the number of mCs divided by numbers of cytosines (Cs) plus thymines (Ts). Pearson’s correlation coefficients were computed between biological replicates of WGBS data (**Fig. S12 & S13**). All pairwise correlation coefficients between biological replicates exceeded 0.8. An average value across biological replicates was used to represent the methylation level of the species. To compare methylation level between genomic regions, we calculated mean methylation level across all methylated cytosines (mCs) in this region.

### ATAC sequencing and ACR identification

The nuclei were extracted from leaves, and the nuclei pellet was resuspended in the Tn5 transposase reaction mix. The transposition reaction was incubated at 37°C for 30 min. Equimolar Adapter1 and Adapter 2 ligation and PCR amplification were performed for library preparation. Libraries were purified with the AMPure beads and quality controlled with Qubit. The clustering of the index-coded samples was performed on a cBot Cluster Generation System using TruSeq PE Cluster Kit v3-cBot-HS (Illumina) according to the manufacturer’s instructions. After cluster generation, all libraries were sequenced on an Illumina Hiseq platform and 150 bp paired-end reads were generated. Nextera adaptor sequences were trimmed from the raw reads using skewer v0.2.2 (Jiang *et al*., 2014). Cleaned reads were then aligned to six reference genomes using Bowtie v2.4.4 (Langmead & Salzberg, 2012) with the following parameters: “bowtie2 --very-sensitive -p 20” (**Table S10**). Aligned reads were sorted using SAMtools (Danecek *et al*., 2021) and clonal duplicates were removed using Picard v2.18.21 (http://broadinstitute.github.io/picard/). Accessible chromatin regions (ACRs) were identified using the “callpeak” function implemented in MACS3 (Zhang *et al*., 2008). Distances between ACRs and adjacent genes were calculated using Homer v5.1 (Heinz *et al*., 2010). The distribution of ACR peak annotation was visualized by Deeptools v.3.5.2 (Ramírez *et al*., 2016).

### Methylation difference in genic region and promoter ACR

The methylation level for each species was assessed following two steps: 1) the methylation level at CG, CHG, and CHH sites was first calculated for each species by averaging across all individuals; 2) the total methylation level for each gene, 3kb up-/down-stream region, or promoter ACR region was calculated by averaging the methylation level of all mCs in the region.

We compared the methylation levels of introgressed and non-introgressed genes. Gene and the 3kb-flanking sequences were partitioned into 20 equidistant bins, respectively. The methylation level of the bin was defined as the mean of methylation ratios of all methylated sites within the bin. The mean methylation levels of all introgressed and non-introgressed genes were taken across all 19 species and compared using Student’s t-test. The 95% confident intervals for two groups of genes were also generated by *t*-distribution.

For genes with ACRs identified in the promoter regions, we also compared the methylation levels of introgressed and non-introgressed ACRs in the promoter regions. Only ACRs at 0 - 3kb upstream of genes were considered and were grouped based on whether the downstream gene was introgressed or not. We partitioned each ACR into ten equidistant bins and partitioned the 3kb upstream/downstream sequences flanking ACRs into 20 bins. The same protocols were used to calculate and compare methylation levels between introgressed and non-introgressed ACRs.

### MCMCGLMM model construction

First, we summarized how introgression affects gene expression level using the following linear model:

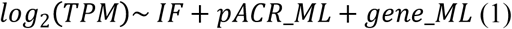

where *TPM* represented Transcript Per Kilobase Million for gene expression level; *IF* represented introgression frequency; p*ACR*_*ML* and *gene*_*ML* represented the methylation level for promoter ACR and gene body, respectively. A Phylogenetic Generalized Linear Mixed model (PGLMM) was applied using R package ‘MCMCglmm’ (Hadfield, 2010). A subtree with 19 species was converted to a variance-covariance matrix and taken as the random effect. Secondly, we evaluated how introgression affected variation of gene expression across the 19 species with the following model:

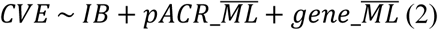

where CVE represented the coefficient of variation of gene expression across 19 species; *pACR_ML* and *gene*_*ML* represented the mean methylation level for promoter ACR and gene body across the 19 species, respectively. The MCMCglmm model was run with parameter set to ‘prior = list(R = list(V = 1, nu = 0.002), G = list(G1 = list(V = 1, nu = 0.002))), nitt = 10000, burnin = 1000, thin = 10’ in both models. Non-significant variables were removed and the best formula was determined by the deviance information criterion (DIC) and shown as in formula (1) and (2), respectively. The marginal and conditional *R*^2^ values were calculated following the methodology proposed by Nakagawa et al. (2017), which decomposes total variance into components explained by fixed effects (marginal *R*^2^) and both fixed and random effects (conditional *R*^2^) in generalized linear mixed models. More materials and methods can be found in **supplementary texts 1.1 – 1.11**.

## Data availability

The raw sequence data reported in this paper have been deposited in the Genome Sequence Archive (Chen *et al*., 2021) in National Genomics Data Center (Bao *et al*., 2024), China National Center for Bioinformation / Beijing Institute of Genomics, Chinese Academy of Sciences with the GSA number CRA029105 for the species resequencing libraries, CRA029292 for WGBS libraries, CRA029333for the RNA-seq libraries and CRA029357 for the ATAC-seq libraries that are publicly accessible at https://ngdc.cncb.ac.cn/gsa. A review link is available (ATAC-seq: https://ngdc.cncb.ac.cn/gsa/s/404Y95Lm; RNA-seq: https://ngdc.cncb.ac.cn/gsa/s/Rh0idJR1; WGBS: https://ngdc.cncb.ac.cn/gsa/s/bsQLe96e; Resequencing: https://ngdc.cncb.ac.cn/gsa/s/15z8IGfX).

## Code availability

All codes used for main analyses in this paper are available for download from https://github.com/furr-rui/protocols-for-widespread_introgression-in-Quercus_species

## Acknowledgments

This project was supported by National Natural Science Foundation of China (32371689) granted to J.C. This research was also supported by National Natural Science Foundation of China (32501371) and Zhejiang Provincial Natural Science Foundation of China (LMS25C030001) to R. F. We appreciated Pan Li, Yang Liu, Lei Cai, Honghua Xu, Xinjie Zhao, Yu Tang for collecting plant samples, and Prof. Fang K. Du for providing an early access to the genome of *Quercus aquifolioides*.

## Author Contributions

J. C. and R. F. designed and managed the project. Y. Zhu and R.F. analyzed the data, with contributions to gene expression analysis from M. Z., Y. Zhu, Y. Li, X. T., J. W., Y. Liu, Y. Zhang, Y. Lin, Z. Z., C.P. and H. W. contributed to plant material collection. Y. Zhu, R. F., C. J., A. K., and M. L. wrote the manuscript. Y. Zhu and R.F. contributed equally.

