## Supplemental texts for "Pervasive historical introgression across oak phylogeny shapes gene expression"

### Supplementary texts

#### 1. Supplementary materials & methods

##### *1.1 Estimating distribution ranges*

The distribution data for the 67 *Quercus* species in this study were obtained from the Global Biodiversity Information Facility (GBIF occurrence was downloaded on March 23rd 2025, https://doi.org/10.15468/dl.84tzej). Occurrence records were filtered according to the native habitat information of each species. Baseline distribution maps for individual species were generated using the R packages 'sf' (40) and 'rnaturalearth'(https://cran.r-project.org/web/packages/rnaturalearth/index.html). To quantify the geographic range of oaks, duplicate occurrence records within each 1° × 1° latitude–longitude grid cell were removed, and the resulting data were standardized using the natural logarithm. We also quantified the number of co-occurring *Quercus* species within all distribution grid cells of each focal species, which was used to represent the extent of range overlap between the target species and other *Quercus* species.

##### *1.2 Quercus super pan-genome construction*

High-quality chromosome-level genomes of nine *Quercus* species were downloaded from public databases, which represent five of eight sections in *Quercus* phylogeny, including *Q. gilva* (section *Cyclobalanopsis*), *Q. acutissima* and *Q. variabilis* (section *Cerris*), *Q. aquifolioides* (section *Ilex*), *Q. rubra* (section *Lobatae*), *Q. lobata*, *Q. dentata*, *Q. mongolica*, and *Q. robur* (section *Quercus*). The *Quercus* super pan-genome was constructed based on nine high-quality genomes. First, genomic synteny between the *Q. variabilis* genome and other eight genomes was evaluated using WGDI (1) and NGenomeSyn (2) with default parameter settings. Orthologous gene families (OGs) were identified using OrthoFinder v2.5.4 (3) with default parameters. All OGs were classified into three groups: 1) the core gene families that were present in all nine genomes; 2) the dispensable gene families that were shared by at least two but not every genome; 3) the species-specific gene families that were present in only one genome. We chose *Q. variabilis* genome as the backbone of the super pan-genome since it has the best assembly quality and integrity of all nine genomes. All other eight genomes were aligned to *Q. variabilis* genome based on Orthofinder results. A graph-based super pan-genome of genus *Quercus* was constructed using the minigraph-cactus pipeline implemented in Cactus v2.5.2 (4). The pipeline was run with the following parameters: 'cactus-pangenome --reference Quercus_variabilis --vcf --giraffe --gfa --gbz'. The *Quercus* super pan-genome graph was saved under multiple graph formats, including VG, GFA, and HAL, in order to enable downstream genetic variant calling.

##### *1.3 Transposable element identification*

A uniform workflow was used to identify and annotate transposable elements (TEs) in all nine genomes. TEs were initially identified using the Extensive *de novo* TE Annotator (EDTA) v2.1.2 pipeline (5), which implemented both well-performing structure- and homology-based identification methods. Subsequently TE outputs were filtered by EDTA with default parameters to gain a high-quality non-redundant TE library and classify TEs into families. The "LTR/unknown" TEs family was further re-classified by DeepTE (6).

To estimate the insertion time of TEs, we extracted full-length LTR retrotransposons (fl-LTRs) and calculated the insertion time using the formula $T = K/2\mu$, where $K$ is the sequence divergence rate between its 5' and 3' LTRs which is estimated from Jukes-Cantor model (7) and $\mu$ is the neutral mutation rate, which equals to $2.69\times10^{-9}$ per base pair per year in *Quercus* (8).

##### *1.4 Recombination rates, gene GC-content calculation*

Based on the genetic map of *Quercus robur* (9), we converted the recombination rates to that of *Quercus variabilis* using the genomic coordinates based on the pan-genome alignments with the following 'halLiftover' command line 'halLiftover Quercus9-pg.full.hal Q_robur Q_robur_geneticmap_sort.bed Q_variabilis Qv_converted_recomb_rate.bed' (10). We then calculated recombination rates in each 10 Kb genomic sliding windows for all chromosomes using in-house Python scripts, as well as the average recombination rate across the gene body. The average GC contents at 1st, 2nd and 3rd positions of amino acid codons (GC1, GC2, GC3) and the overall GC content averaged for each protein coding gene were also calculated using in-house Python scripts.

##### *1.5 Divergence time and diversification rate estimation*

Based on the phylogenetic tree, divergence time was estimated by MCMCTree v4.10 in the PAML v4.10 package (11) with 'HKY85' nucleotide substitution model and an independent molecular clock rate for each branch. The same five fossil records as in (12) were used to recalibrate the branch length (**Table S4**). The MCMC chains were first run for 3 million iterations as burn-in, and then were sampled every 500 iterations until a total of 50,000 samples were collected (i.e., 25 million iterations). Tracer v1.7.2 (13) was used to confirm the MCMC convergence of each run by ensuring the effective sample size of all parameters greater than 200. For both individual-based and species-based phylogenetic trees, three independent MCMCTree runs were generated with different random number seeds and very similar results were obtained for both phylogenetic trees.

We used the Bayesian Analysis of Macroevolutionary Mixture (BAMM) v2.5.0 (14) software to estimate the diversification rate of the *Quercus* genus. The recalibrated phylogenetic tree with divergence time estimated by MCMCtree was used as an input. To account for incomplete taxon sampling, we calculated the sampling fraction for each section of genus *Quercus* based on the number of species reported in (12). Absent taxa were added to randomly selected positions in corresponding lineages (**Table S5**). The BAMM analyses were run for 10 million iterations, and sampled every 1,000 iterations. The first 30% samples were discarded as burn-in, and the remaining samples were summarized and plotted using BAMMtools v2.5.0 (15).

##### *1.6 Test for identity-by-descent*

We performed IBD analyses based on all 9,795,402 SNPs. Beagle v5.4 (16) was used to phase and impute SNPs. Subsequently, Truffle v1.38 (17) was employed for IBD analyses with the parameters '--mindist 2000 –segments'. The results were then plotted in R.

##### *1.7 The correlation tests between introgression and genetic divergence*

We then tested how introgression changed with species divergence by regressing $f_d$ on either Nei's $D_{xy}$ or the phylogenetic distance between two species. Nei's $D_{xy}$ was calculated across 100-kb sliding windows (50 kb step) using 'popgenWindows.py' (18). The phylogenetic distance was calculated using 'patristicdistmatrix' function in phytools (19) by summing over the lengths of all branches connecting the two species that are tested. Mean $f_d$ between P2 and P3 was taken across all possible triplets containing P2 and P3. A Mantel test was used to assess the significance of the regression of $f_d$ on either $D_{xy}$ or phylogenetic distance.

##### *1.8 Gene ontology and plant ontology analyses*

Gene ontology (GO) enrichment analysis was performed for selected candidate genes using *Arabidopsis thaliana* orthologues (BLASTP with e-value < 1e-5) in the PANTHER Classification System on the Gene Ontology Database (released 17 January 2024, http://geneontology.org/) (20). Fisher's exact test was used to determine the enrichment significance of GO terms, and *p*-values were further adjusted by FDR to correct for multiple testing. Plant ontology (PO) database was downloaded from Plant ontology consortium (21) (https://archive.plantontology.org). A hypergeometric test was used to calculate *p*-values of enrichment significance for both PO anatomy and temporal terms and *p*-values were further adjusted by FDR to account for multiple testing.

##### *1.9 Gene pleiotropy analysis*

We used the number of protein-protein interaction to evaluate the pleiotropic effects for the introgressed and non-introgressed genes. First, we searched candidate gene sequences against *Arabidopsis thaliana* TAIR10 gene database using BLASTP with an e-value cutoff $\leq 1\times10^{-5}$. Secondly, a Protein-Protein Interaction (PPI) network was reconstructed using the STRING database (22) and was visualized by software Cytoscape (23). Finally, the pleiotropic degree for each gene was counted as the number of proteins it interacts with in the network using CytoHubba (23).

##### *1.10 Estimation of $K_a/K_s$ ratios*

SNPs were phased and imputed by Beagle v5.4 (16) using default parameters. To gain species-specific protein coding sequences, SNPs were used to replace the nucleotide at the corresponding positions of the reference CDS sequences using 'vcf2fasta.py' script (https://github.com/santiagosnchez/vcf2fasta). The program 'codeml' was used to estimate $K_a/K_s$ ratio for each gene and each species with two models: 1) the null model that assumes a single $K_a/K_s$ ratio for all branches, and 2) the free-ratio model that assumes an independent $K_a/K_s$ ratio for each branches (11). The likelihood ratio test was used for model selection. Genes with significantly better performance in the free-ratio model and with $K_a/K_s$ ratio > 1 in more than 50% oak species were considered as candidate genes under positive selection in genus *Quercus*.

##### *1.11 Prediction of transcription factors and their regulated genes*

To gain deeper insights into the difference in regulatory mechanisms between introgressed genes and non-introgressed genes, we utilized the online tool PlantTFDB v5.0 (https://planttfdb.gao-lab.org/prediction.php) (24) to predict transcription factors based on the best hits in *Arabidopsis thaliana*, using annotated genes of *Quercus variabilis* ($p$-value $\leq 0.01$). Promoter regions extending 3000 bp upstream of each gene were extracted and predicted as regulated genes using the PlantRegMap (https://plantregmap.gao-lab.org/binding_site_prediction.php) (24), applying a $p$-value threshold of $\leq 1e-7$. The resulting transcription factor-regulated genes were then summarized and visualized using R v4.3.1.

#### 3. Supplementary figures

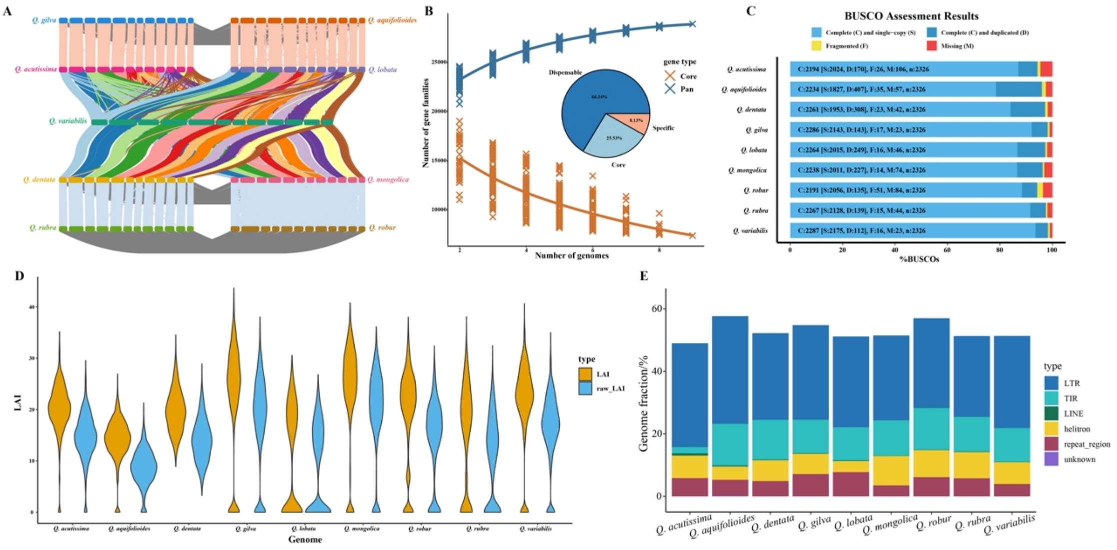


**Figure S1 Genomic features of nine oak reference genomes used for constructing the super-pan genome of the genus *Quercus*. (A)** The collinearity map of reference genomes of nine oak species. Chromosomes are represented by colored blocks. **(B)** Accumulation curves of the super pan-genome and core-genome. The scatter plot shows the number of gene families present at different frequencies across the nine reference genomes; the pie chart illustrates the distribution of gene family categories. Solid lines represent the median number of gene families. **(C)** BUSCO assessment results of nine reference genomes. **(D)** LTR Assembly Index (LAI) distribution. Blue violin plots represent raw LAI values, indicating the proportion of intact LTR retrotransposons in sliding windows; yellow violin plots represent corrected LAI values. **(E)** Proportion of repeat sequences in *Quercus* genomes.

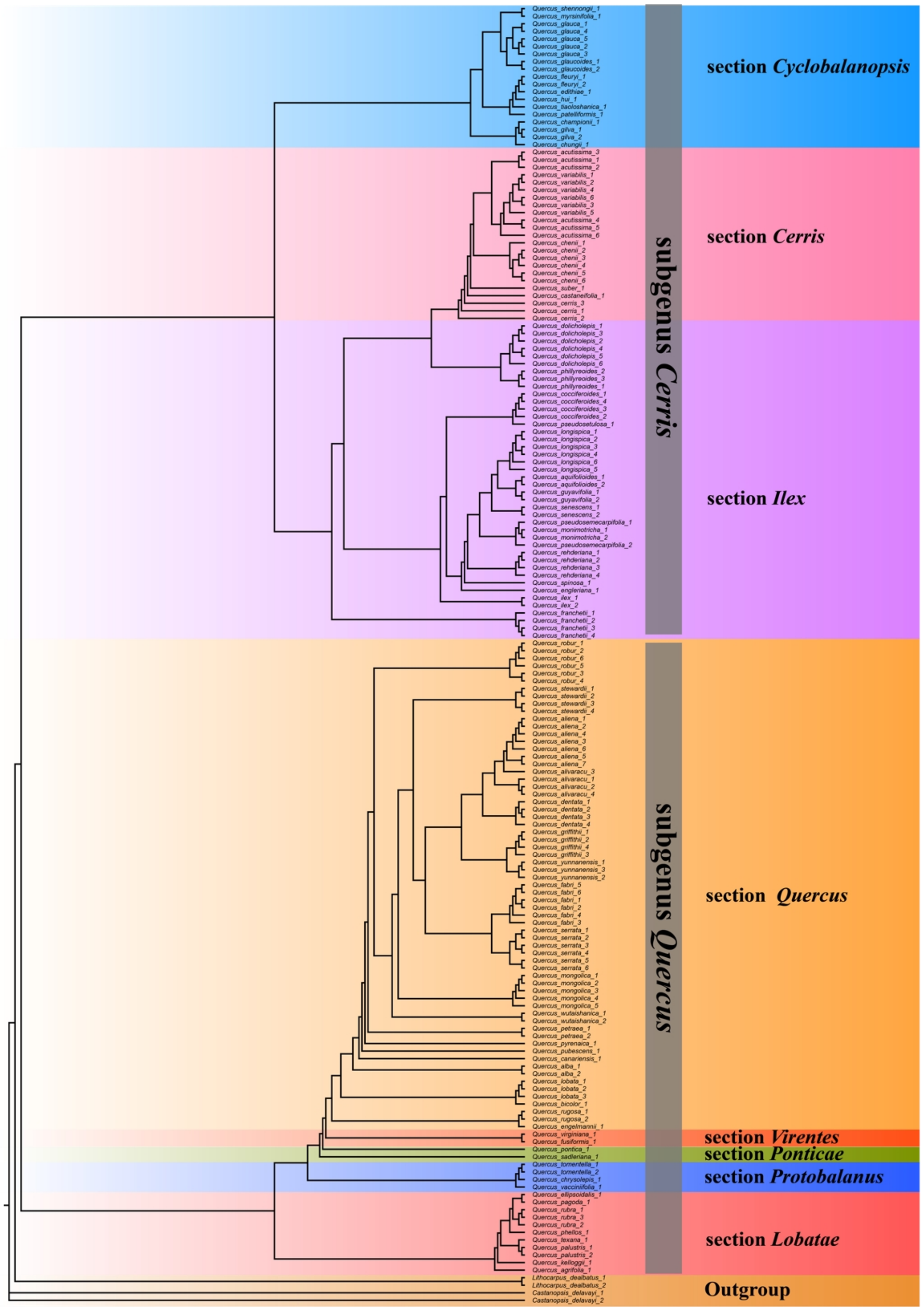


**Figure S2 The phylogenetic tree of genus *Quercus* based on unlinked SNPs across genomes of 168 individuals from 67 species.**

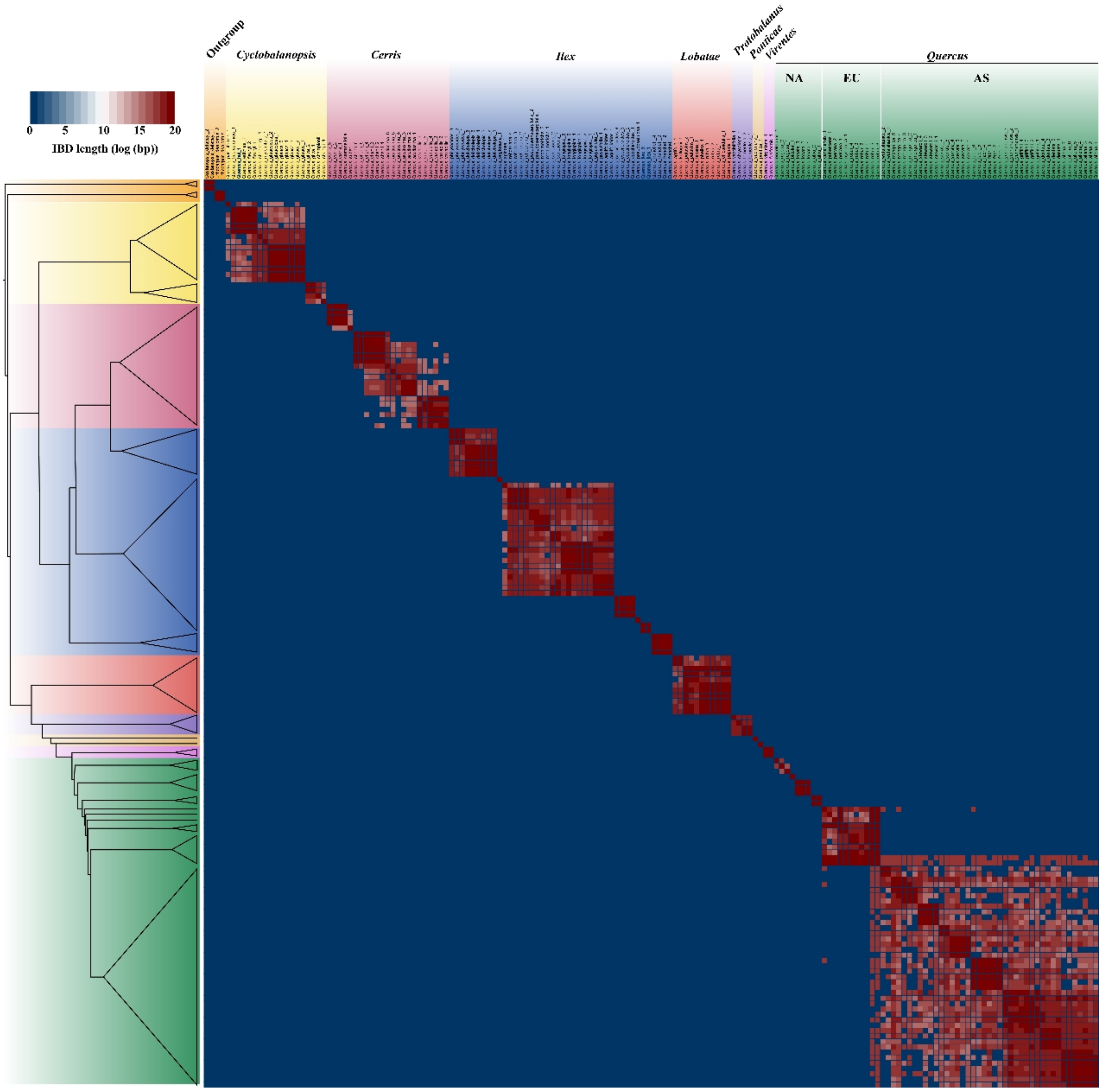


**Figure S3 Shared identity-by-descent (IBD) blocks between *Quercus* species.** Heatmap indicating the total length of IBD blocks for each pair of comparisons.

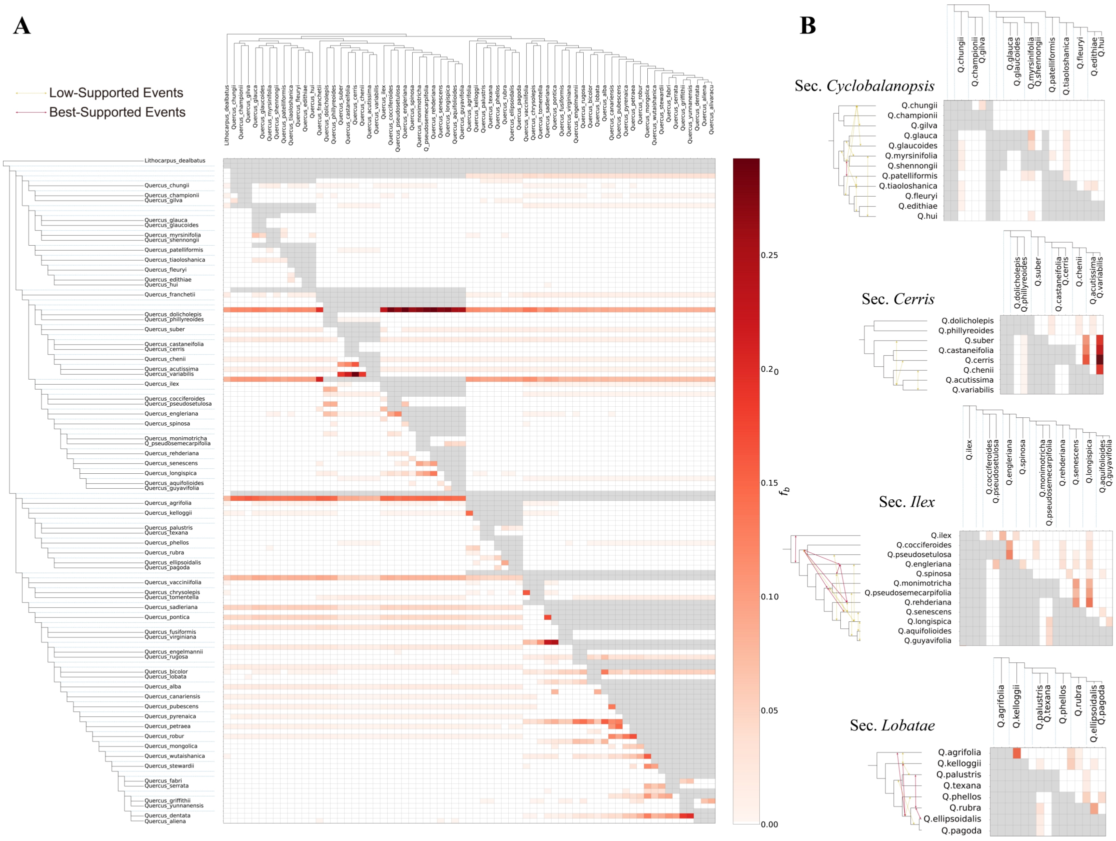


**Figure S4 *f*-branch results and inference of introgression events. (A)** *f*-branch results of 67 *Quercus* species. **(B)** *f*-branch results and inference of introgression events in 4 main Sections. Red arrows indicate best-supported introgression events that passed both DCT/BLT tests, and yellow arrows indicate low-supported introgression events.

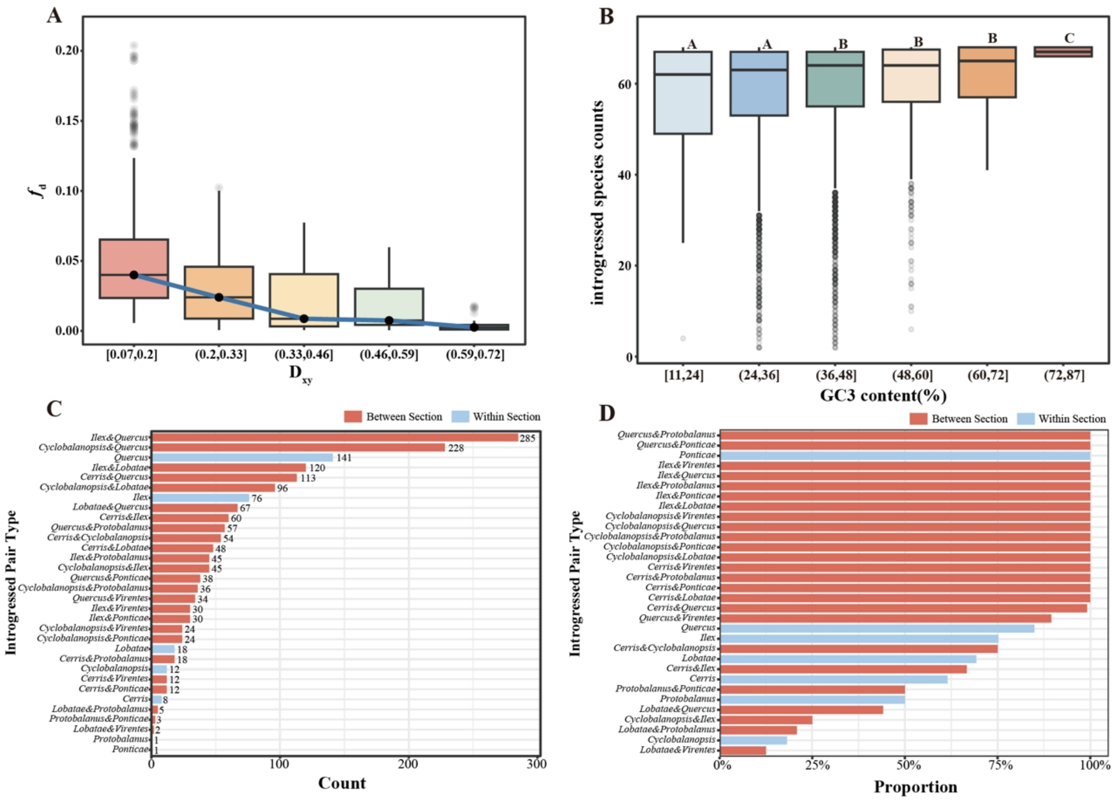


**Figure S5 Correlation between introgression frequency and strength with phylogeny among *Quercus* species. (A)** Correlation map of introgression strength and genetic distances ($D_{xy}$) among *Quercus* species. The horizontal coordinate represents $D_{xy}$ interval. **(B)** Correlation map of introgression species counts and GC3 content interval among *Quercus* plants. The horizontal coordinate represents GC3 content interval; **(C)** Total number of species pairs within and between sections that significant introgression signals were identified in. **(D)** Proportion of introgressed species pairs in **(C)** relative to all possible species pairs within and between sections. Blue bars represent the proportion within sections, and red bars represent the proportion between sections.

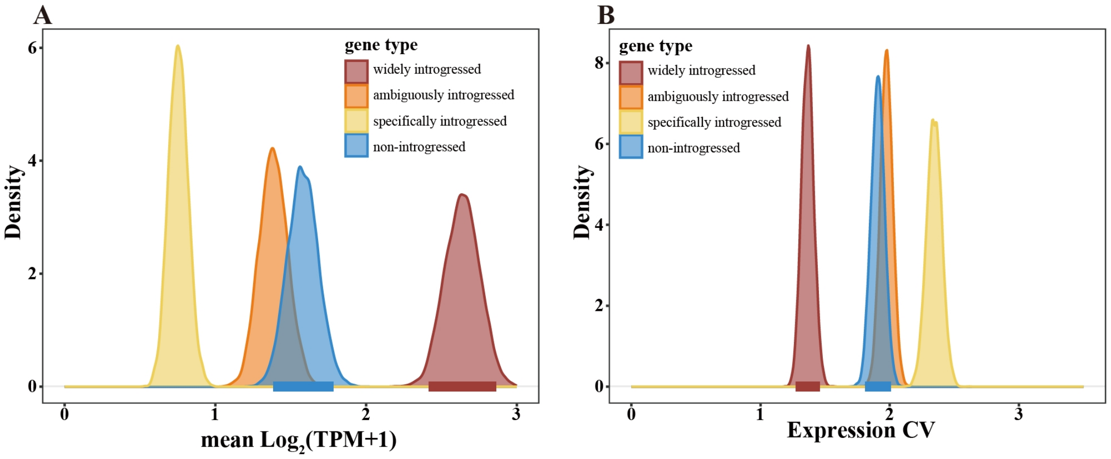


**Figure S6 The distributions of gene expression level and coefficient of variation (CV) for four types of genes based on introgression frequency. (A)** Distributions of mean expression level averaged across 19 oaks that were generated by bootstrapping each for 1000 times. **(B)** Distributions of coefficient of variation (CV) of gene expression across 19 oaks that were generated by bootstrapping each for 1000 times.

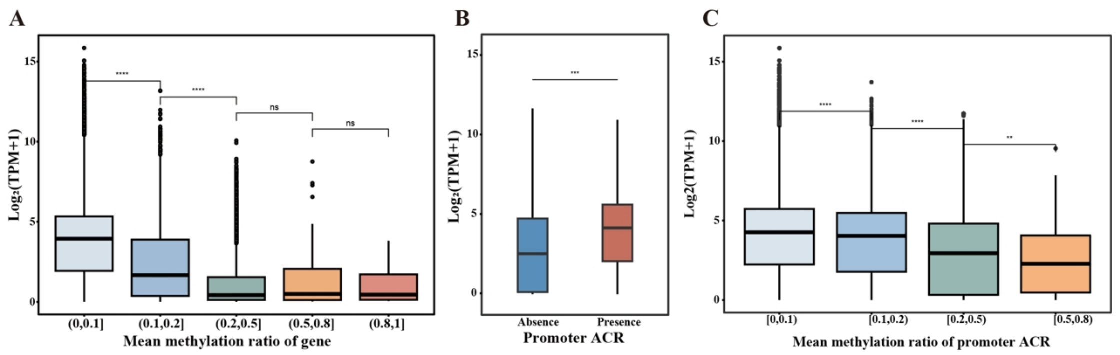


**Figure S7 The relationship between gene expression level and gene methylation level (A), the presence or absence of promoter ACR (B), or the methylation level of promoter ACR (C).**

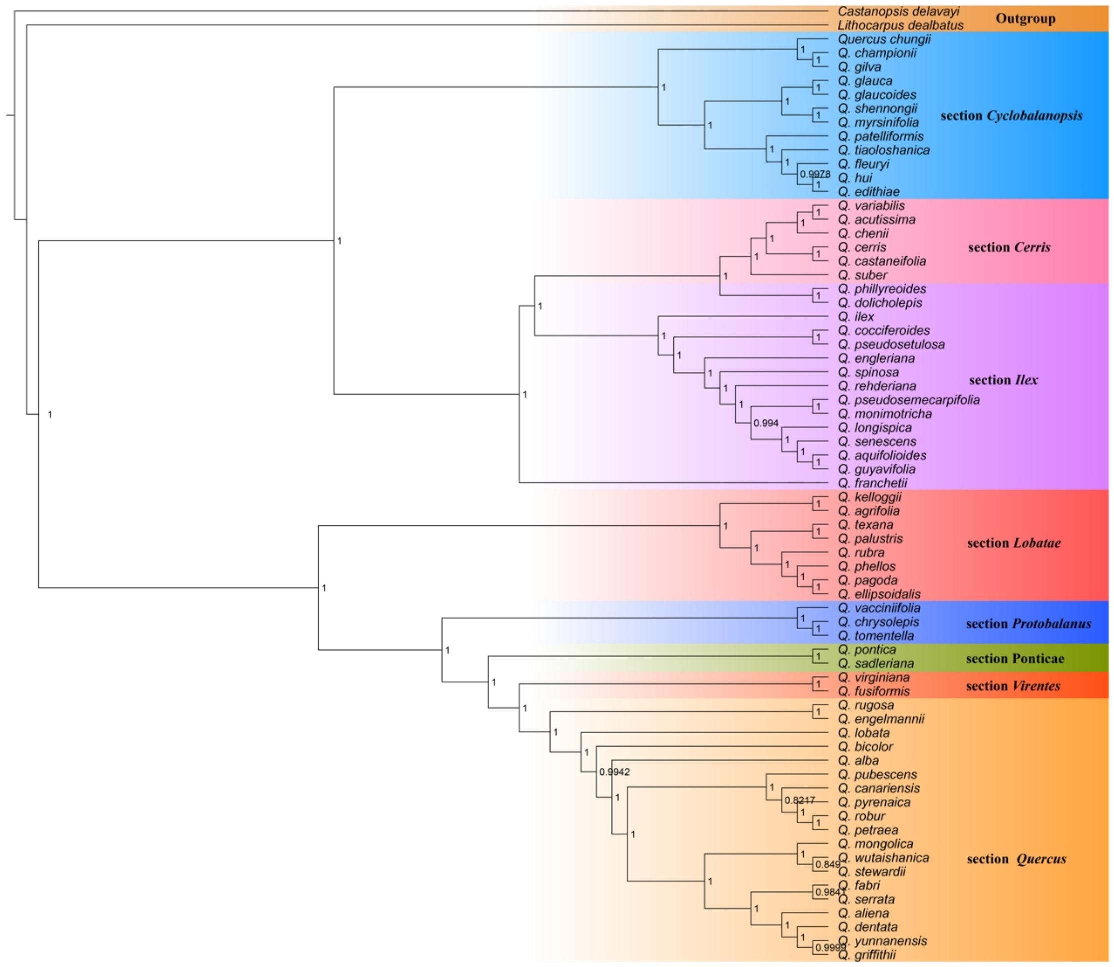


**Figure S8 Species tree of *Quercus* inferred by ASTRAL.** The species tree was reconstructed using ASTRAL v1.16.3.4 based on 9,804 individual gene trees. Numbers at internal nodes indicate local posterior probabilities (LPP).

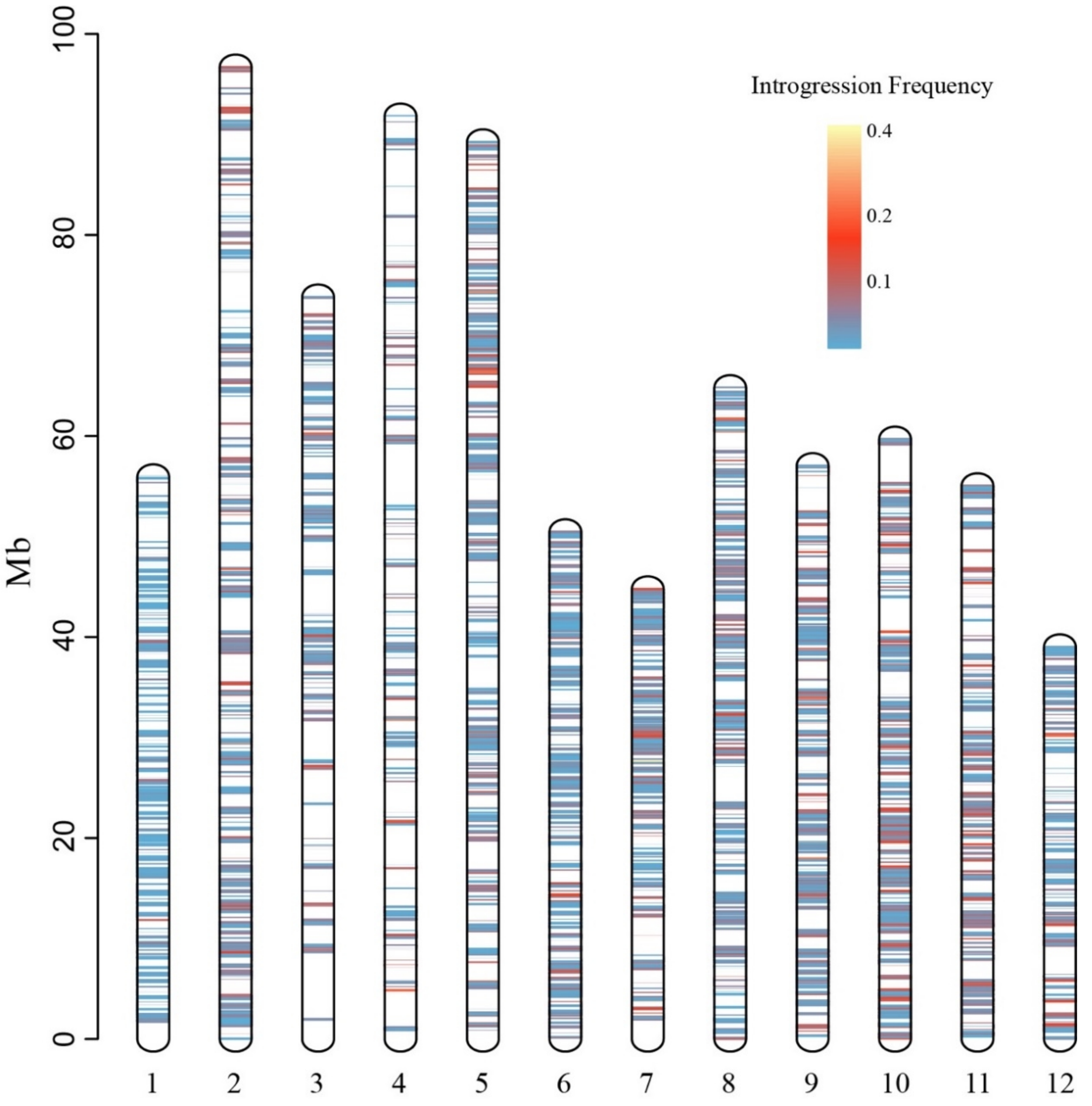


**Figure S9 The genomic positions of all introgressed genes identified in species pairs across 67 oaks were mapped to the genome of *Q. variabilis*.** Colors represent the introgression frequency.

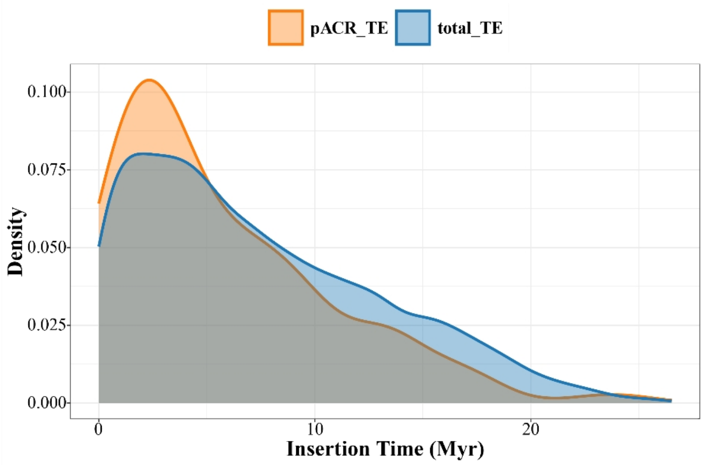


**Figure S10 The distributions of insertion time for all TEs across the genome (blue) and TEs only found promoter ACRs (orange).**

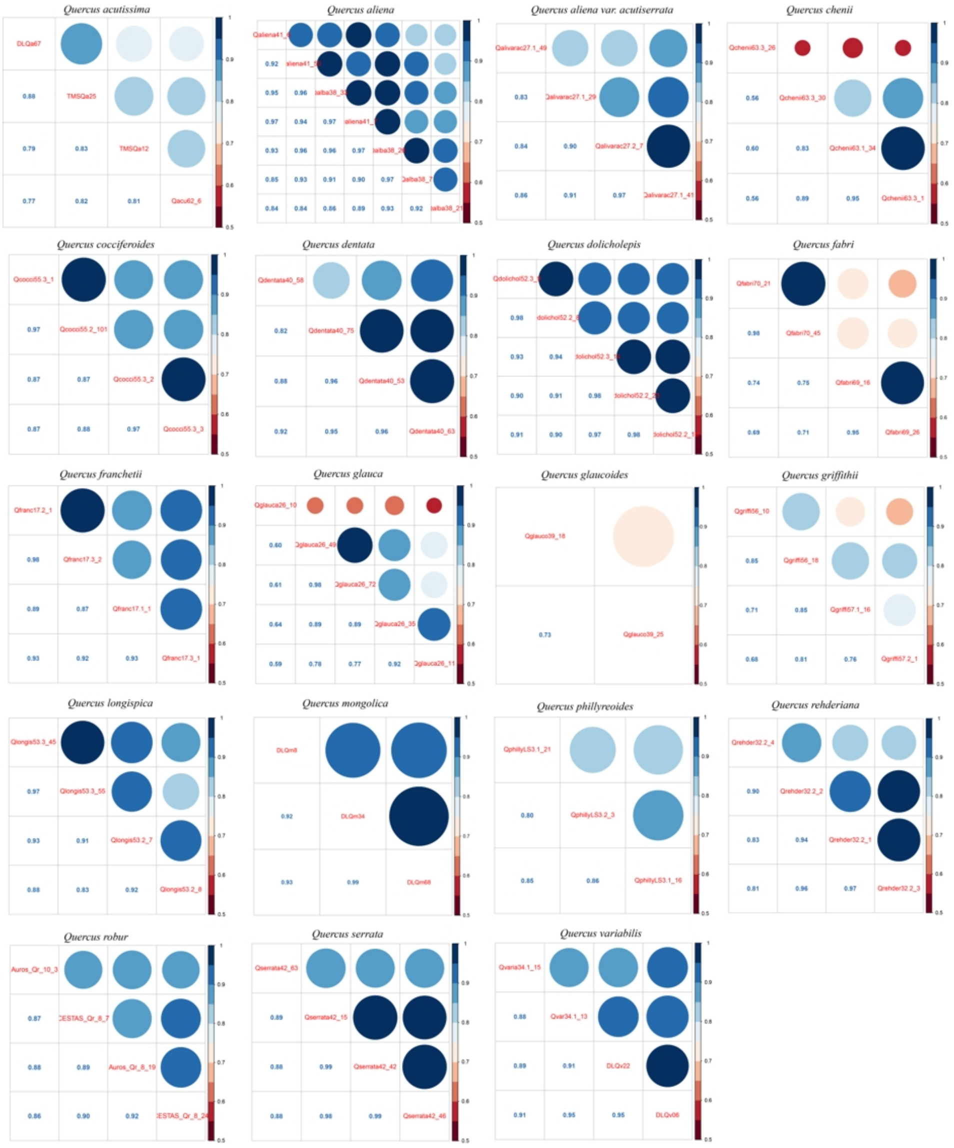


**Figure S11 The correlation coefficient matrices based on gene expression difference between biological replicates for transcriptomic sequencing (RNA seq) for 19 *Quercus* species.** Both color and dot size represent the correlation coefficient, which is also shown in the lower diagonal.

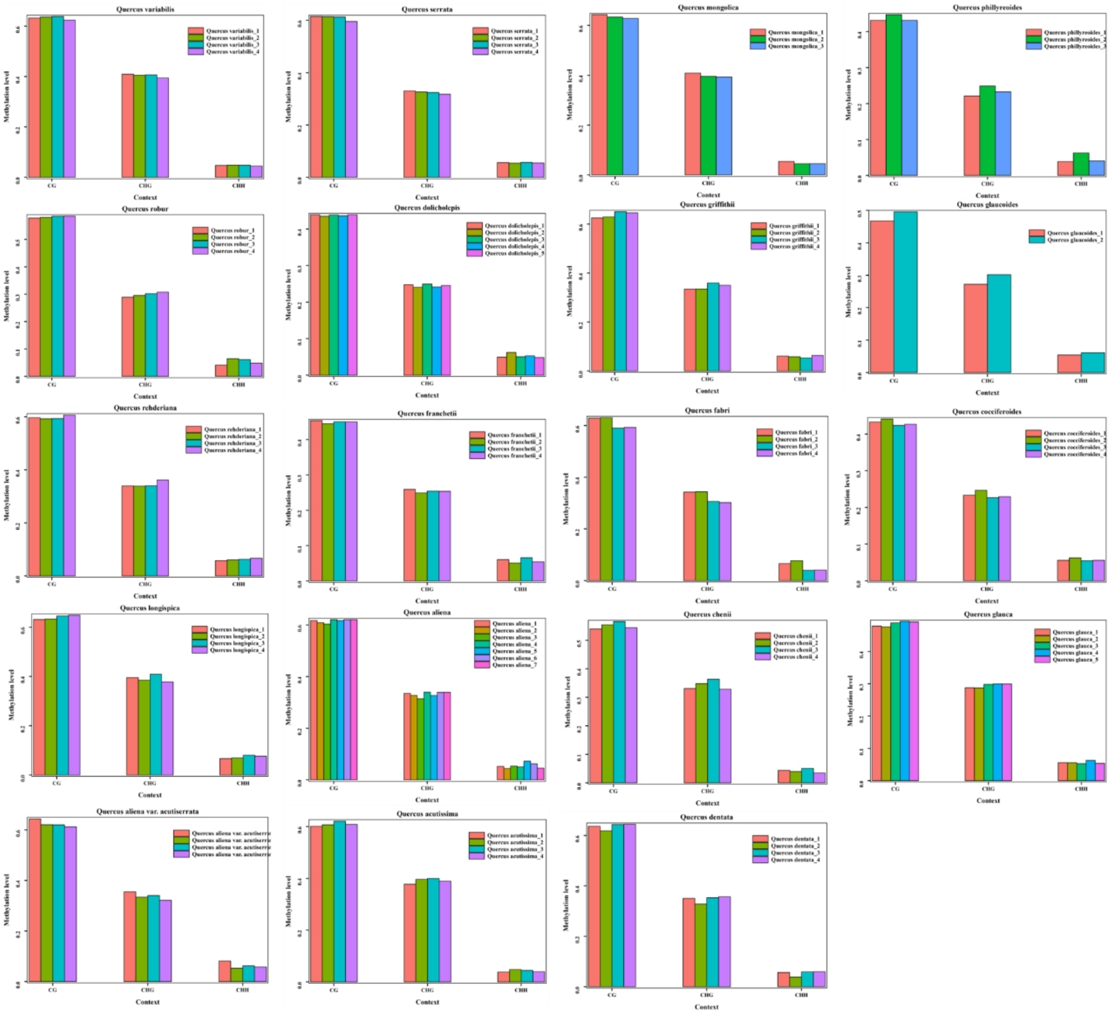


**Figure S12 The mean methylation levels for 19 species at three different types of sites (CG, CHG, CHH) across the genome in different individuals.**

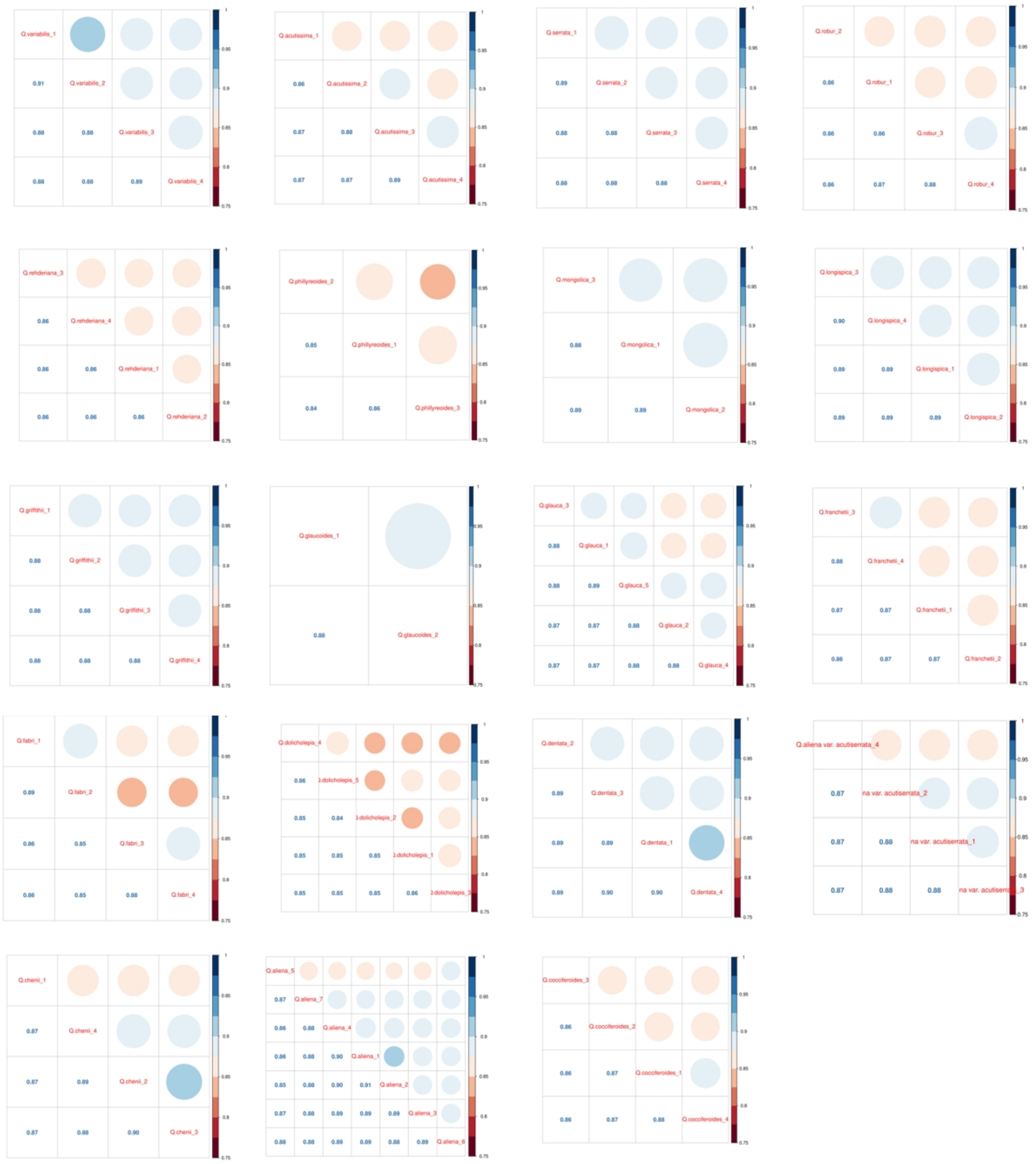

**Figure S13 The correlation matrices based on methylation difference between biological replicates for methylation sequencing (WGBS) for 19 *Quercus* species**. Both color and dot size represent the correlation coefficient, which is also shown in the lower diagonal.
